# Structure and dynamics of semaglutide and taspoglutide bound GLP-1R-Gs complexes

**DOI:** 10.1101/2021.01.12.426449

**Authors:** Xin Zhang, Matthew J. Belousoff, Yi-Lynn Liang, Radostin Danev, Patrick M. Sexton, Denise Wootten

## Abstract

The glucagon-like peptide-1 receptor (GLP-1R) regulates insulin secretion, carbohydrate metabolism and appetite, and is an important target for treatment of type II diabetes and obesity. Multiple GLP-1R agonists have entered into clinical trials, such as semaglutide, progressing to approval. Others, including taspoglutide, failed through high incidence of side-effects or insufficient efficacy. GLP-1R agonists have a broad spectrum of signalling profiles. However, molecular understanding is limited by a lack of structural information on how different GLP-1R agonists engage with the GLP-1R. In this study, we determined cryo-electron microscopy (cryo-EM) structures of GLP-1R-Gs protein complexes bound with semaglutide and taspoglutide. These revealed similar peptide binding modes to that previously observed for GLP-1. However, 3D variability analysis of the cryo-EM micrographs revealed different motions within the bound peptides and the receptor relative to when GLP-1 is bound. This work provides novel insights into the molecular determinants of peptide engagement with the GLP-1R.

## INTRODUCTION

G protein-coupled receptors (GPCRs) mediate diverse physiological processes through selective modulation of cellular function and thus represent important drug targets (Hauser et al. 2017). The glucagon-like peptide-1 receptor (GLP-1R), a class B1 subfamily GPCR, signals primarily through the stimulatory G (Gs) protein. It mediates important physiological effects of the endogenous peptide, GLP-1, in regulating insulin secretion, carbohydrate metabolism and appetite, and thus is well established as a long term clinical target for treating type II diabetes and obesity (Graaf et al. 2016). Multiple GLP-1R peptide agonists that are used clinically or that have entered clinical trials display differential signalling profiles and different clinical efficacies or side-effect profiles (Graaf et al. 2016, Sun et al. 2014, Koole et al. 2010, Jones et al. 2018, Fletcher et al. 2018, Pickford et al. 2020). Of note are two GLP-1 derived peptides, semaglutide and taspoglutide that have distinct clinical profiles. Once-weekly semaglutide, approved by the U.S. Food and Drug Administration (FDA) in December 2017, provided superior glucose correction and weight loss in phase 3 clinical trials relative to the two most commonly used therapeutics exenatide and liraglutide (Ahmann et al. 2018, O’Neil et al. 2018). In contrast, taspoglutide, while meeting its primary efficacy end point, failed in late phase 3 clinical trials due to intolerable side effects (Rosenstock et al. 2013). Notably, both peptides are analogues of human GLP-1 (7-36); semaglutide has two amino acid substitutions (Aib8, Arg34) and is derivatised at Lys26 with C18 diacids via a γGlu-2xOEG linker, while taspoglutide is an Aib8 and Aib35 substituted analogue (Figure S1A). Their high sequence identity but distinctive clinical profiles make understanding the molecular details that underlie the differences in their mechanism of action important for future development of novel drugs targeting GLP-1R.

Advances in cryo-electron microscopy (cryo-EM) have enabled structure determination of GPCRs coupled to heterotrimeric G proteins (Garcia-Nafria and Tate 2019), with recently solved cryo-EM structures of class B1 GPCRs achieving global resolutions of 2.5 Å or below (Liang, Belousoff, Fletcher, et al. 2020, Dong et al. 2020, Zhang et al. 2020). Cryo-EM can also capture ensembles of conformations present during vitrification, providing insight into the dynamic behaviour of proteins (Lau et al. 2018, Murata and Wolf 2018), a crucial factor in molecular understanding of agonist binding and GPCR activation. With large, high-resolution (sub-3 Å) data sets, conformational heterogeneity can be parsed into principal components by 3D variability analysis implemented in software packages such as cryoSPARC (Punjani et al. 2017, Punjani and Fleet 2020) to gain insight into relative dynamics of specific ligand-GPCR complexes (Dong et al. 2020, Liang, Belousoff, Fletcher, et al. 2020, Zhang et al. 2020). In this study, we report the structures of GLP-1R-Gs complexes with semaglutide and taspoglutide at global resolutions of 2.5 Å. 3D variability analysis was used to reveal differences in the conformational dynamics of semaglutide and taspolutide-coupled GLP-1R-Gs complexes. The consensus and dynamic results were compared with the endogenous peptide GLP1-GLP-1R complex structure reported previously (Zhang et al. 2020), providing new insights into engagement of the GLP-1R by distinct peptide agonists.

## RESULTS

### Structure determination

Human GLP-1R, dominant negative Gαs (DNGαs) (Liang, Zhao, et al. 2018), Gβ1 and Gγ2 constructs were co-expressed in *Trichoplusia ni* (Tni) insect cells to facilitate G protein-coupled complex formation. Formation of GLP-1R-Gs complexes was initiated by addition of 10 μM of either semaglutide or taspoglutide. Stabilization of the complex was achieved by combination of apyrase to remove guanine nucleotides, nanobody 35 (Nb35) that bridges the Gα-Gβ interface (Rasmussen et al. 2011) and use of DNGαs (Liang, Zhao, et al. 2018). Purified semaglutide- and taspoglutide-GLP-1R-Gs complexes were resolved as monodisperse peaks through size exclusion chromatography (SEC) that contained all expected components as confirmed by western blot and Coomassie stained SDS-PAGE (Figures S1B and S1C). Samples were vitrified and imaged on a 300kV Titan Krios cryo-EM, generating particles that exhibited high-resolution features following 2D classification (Figures S1B, S1C and S2). These data were subsequently refined in 3D to yield consensus maps with global resolutions of 2.5 Å for both peptide-bound complexes at gold standard FSC 0.143 (Figures 1, S1B-C and S2). As there was only limited density for the α-helical domain (AHD) of the Gα subunit, this was masked out during the consensus map refinement.

**Figure 1.**
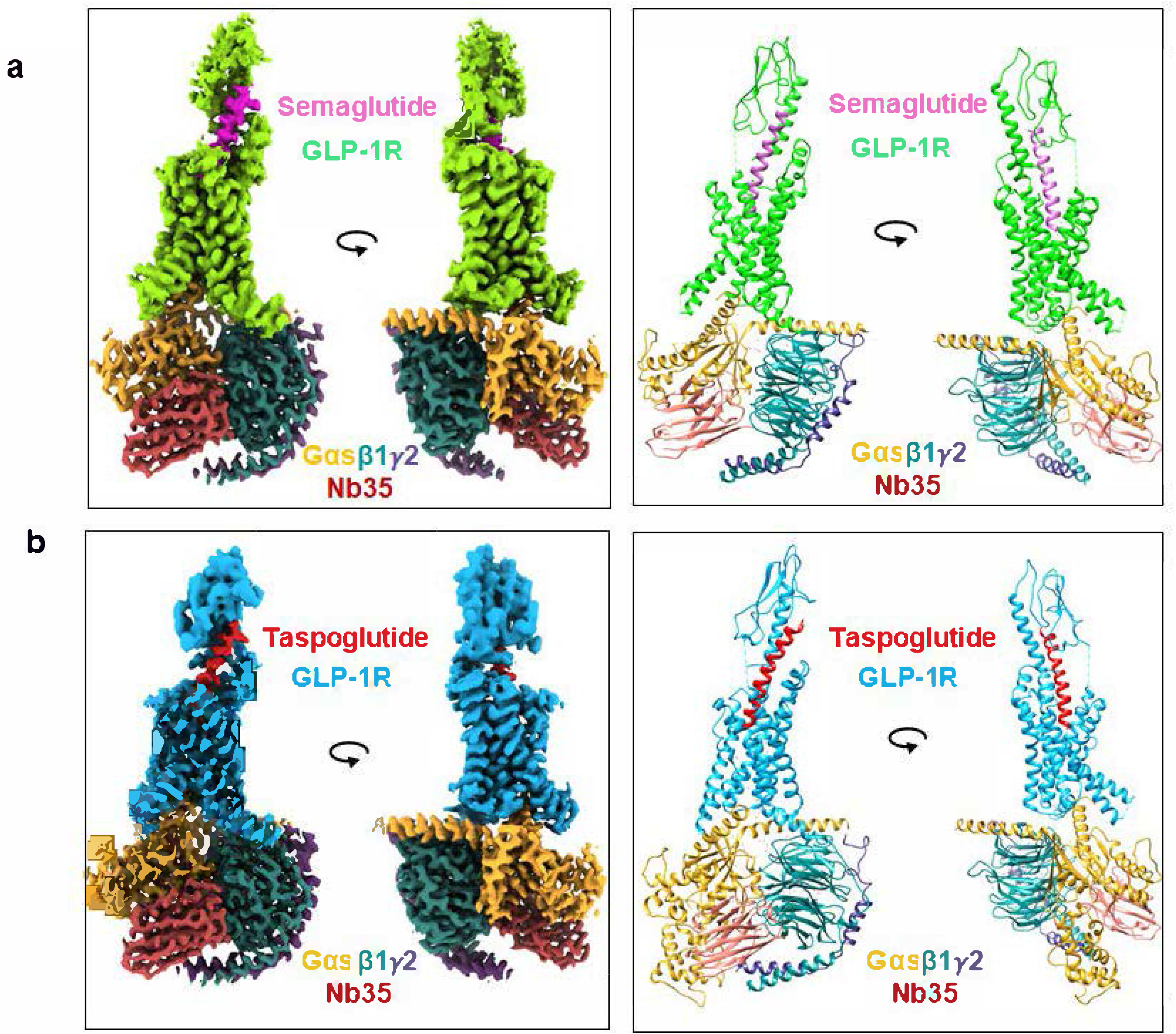
Cryo-EM structures of GLP-1R-G_s_ complexes. **a**, Semaglutide-bound complex; **b**, Taspoglutide-bound complex. Left, Orthogonal views of the cryo-EM maps generated from consensus maps via zone of 2 Å and mask on each component of complex models using UCSF ChimeraX; Right, the models built into the maps, displayed in ribbon format. Colouring denotes the protein segments as highlighted on the figure panels. *Also see Figures S1, S2, S3 and Table S1*.

In the consensus maps the local resolution ranged from 2.4 Å to > 4.0 Å for both complexes (Figures S1B-C) with the highest resolution observed for the G protein, receptor transmembrane domain (TMD) and peptide N-terminus. Lower resolution was observed for the extracellular and intracellular loops, extracellular N-terminal domain (ECD) and peptide C-terminus that is likely due to greater flexibility in these domains. To improve the resolution of poorly resolved regions, focused 3D refinements were performed on the extracellular portion of the TMD and the ECD, for both receptors, which dramatically improved resolution of the ECD, extracellular loops (ECLs) and peptide C-terminus. This enabled robust modelling of the majority of the complex, including confident assignment of most sidechain rotamers (Figure S3). However, residues K130-S136 of the TM1-ECD linker and M340-K342/T343 of ICL3 remained poorly resolved and were omitted from final models. Likewise, there was no density for the receptor C-terminus beyond amino acid L422^8.63^ in the C-terminal Helix 8 (H8), or for residues 24-28 within the far N-terminus of the mature protein, and these were not modelled. While density for the Gαs AHD was poor for both complexes, a higher signal was observed in the taspoglutide data. As such, an additional focused refinement was performed on the G protein of this complex. This improved the resolution of the AHD enabling the backbone of this region to be modelled, although there was insufficient density to robustly model sidechains. In this map, the Gαs AHD was in a relatively closed orientation (Figures 1B and S1Cv).

### Peptide binding to GLP-1R

Similar to most peptide agonists present in active class B1 GPCR complex structures (Zhao et al. 2019, Dong et al. 2020, Liang, Belousoff, Zhao, et al. 2020, Qiao et al. 2020, Liang, Khoshouei, Glukhova, et al. 2018), both semaglutide and taspoglutide adopted a continuous helix with their C-terminus bound to the ECD and N-terminus inserted deeply into the TM core (Figures 1 and S4). The N terminus of GLP-1, particularly residues at position 7, 8, 9, 10, 12, 13 and 15, as well as several residues at the C terminus are crucial for GLP-1 binding and/or activity (Adelhorst et al. 1994). Given the high sequence similarity between GLP-1 and GLP-1 analogues (Figure 1A), it is perhaps not surprising that the packing of the receptor TM core, and the interactions made by the N-terminal half of the peptide and the receptor core are highly conserved in the consensus structures (Zhang et al. 2020) (Figures 2 and S4).

**Figure 2.**
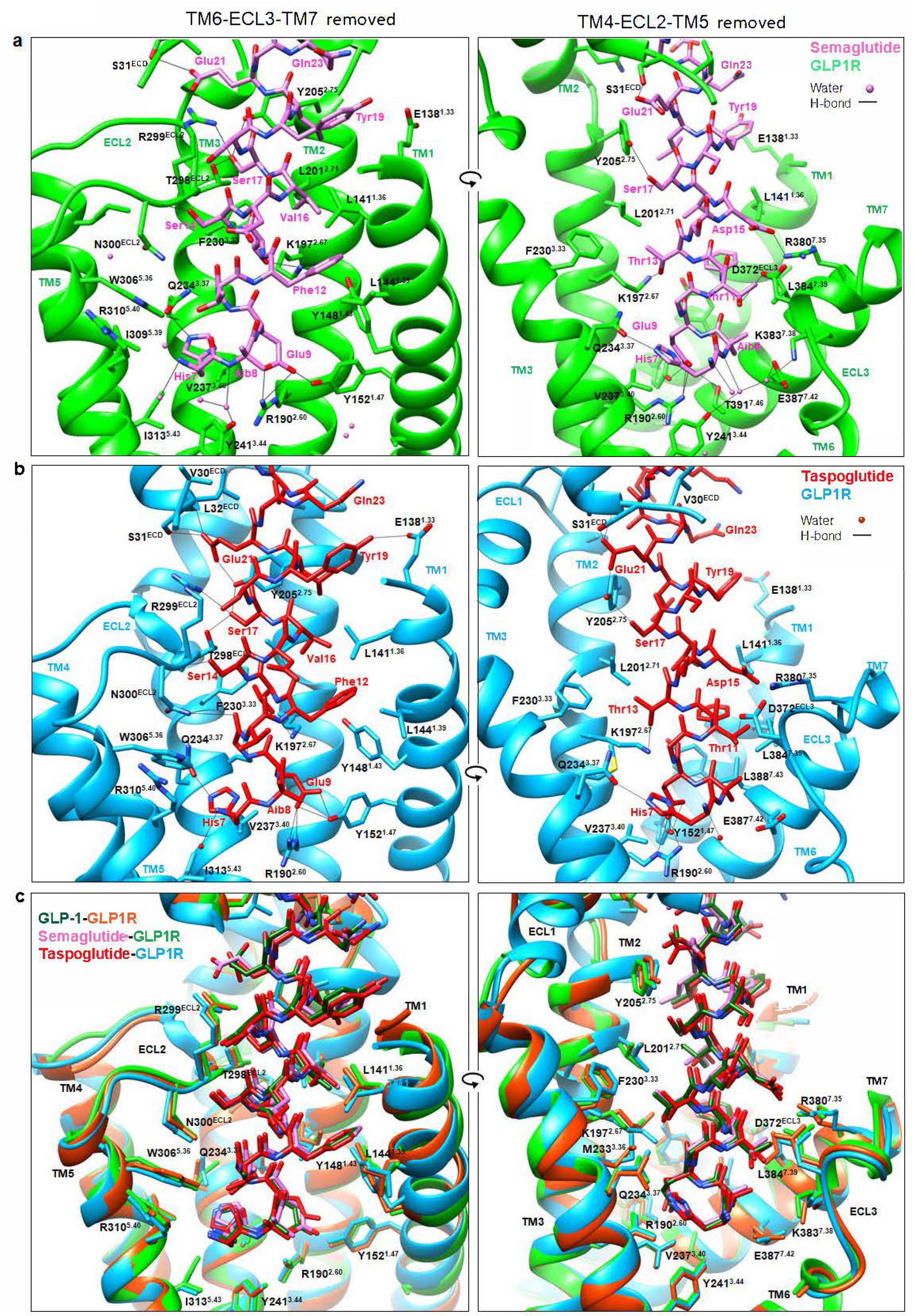
Interactions of semaglutide and taspoglutide within the TM binding cavity of the GLP-1R. GLP-1R residues that interact with peptide or waters (balls) within the GLP-1R binding cavity are displayed in stick format coloured by heteroatom, with the backbone of the receptor in ribbon format. **a**, GLP-1R and semaglutide; **b**, GLP-1R and taspoglutide; **c**, Overlay of GLP-1 (PDB: 6×18), semaglutide and taspoglutide binding sites. Colours are highlighted on the figure panels. For each binding site two views are depicted for clarity: Left, side view of the TM bundle viewed from the upper portion of TM6/TM7 where TM6-ECL3-TM7 have been removed; Right, side view of the TM bundle viewed from the upper portion of TM4/TM5 where TM4-ECL2-TM5 have been removed. Black lines depict hydrogen bonds as determined using UCSF Chimera. *Also see Figures S3, S4 and Table S2*.

Similar to GLP-1, His7 of semaglutide and taspoglutide formed extensive interactions with Q234^3.37^, V237^3.40^, W306^5.36^, I309^5.39^, R310^5.40^ and I313^5.43^ of GLP-1R via Van der Waals forces, and a hydrogen bond with Q234^3.37^; a highly conserved interaction site for class B1 peptides (Liang, Belousoff, Zhao, et al. 2020). Ala8 of GLP-1 is substituted with aminoisobutyric acid (Aib) in semaglutide and taspoglutide, engendering DPP-IV resistance while maintaining high GLP-1R activity (Deacon et al. 1998). In both structures, Aib8 formed hydrophobic interactions with L384^7.39^ and E387^7.42^ and this differed from the predominant interactions of Ala8 of GLP-1 that were with E387^7.42^ and L388^7.43^ (Figure 2). Consequently, alanine mutation of L384^7.39^ had a greater impact on semaglutide- and tasopglutide-mediated cAMP production compared to GLP-1 (Figure S5). The backbone of Aib8 in semaglutide was involved in water-mediated hydrogen bond interactions with Y241^3.44^, K383^7.38^ and E387^7.42^. This network was also observed for the backbone of Ala8 in GLP-1 (Zhang et al. 2020). Although density for the water molecules in this network was not observed in the taspoglutide- bound structure, the almost identical rotamer placements of the three polar receptor residue sidechains suggested that similar interactions were likely to occur (Figure 2C). Glu9 formed hydrogen bonds with Y152^1.47^ and R190^2.60^, and Thr11 formed a hydrogen bond with D372^ECL3^ in both structures (Figures 2A-B). Taken together, His7, Aib8, Glu9 and Thr11 of semaglutide and taspoglutide appear to have similar patterns of receptor interaction to that of GLP-1 that stabilize the peptide above the crucial central polar network (Wootten et al. 2013, Wootten, Reynolds, Koole, et al. 2016) of the TM core that is important for receptor activation.

As GLP-1 derived peptides, semaglutide and taspoglutide have multiple additional polar residues within the N-terminal helix that interact with the receptor core, and participate in conserved hydrogen bond interactions for both peptides. Thr13 interacted with K197^2.67^, while Asp15 interacted with R380^7.35^ at the top of the TM7/ECL3 interface, which provides important coordination of the peptide with these domains as evidenced by mutation of either receptor residue leading to substantial loss of peptide activity (Figure S5). ECL2 that is a key receptor domain involved in G protein coupling (Koole et al. 2012, Dods and Donnelly 2015, Wootten, Reynolds, Smith, et al. 2016, Yang et al. 2016), made a series of coordinated hydrogen bonds with Ser14 (N300^ECL2^), and Ser17 (T298^ECL2^, R299^ECL2^), with the latter serine also involved in a hydrogen bond interaction with Y205^2.75^ at the top of TM2 at the junction with ECL1. Glu21 formed hydrogen bonds with R299^ECL2^ and with the far N-terminus of the receptor ECD to provide coordination of these domains, while Glu21 in taspoglutide also formed a polar interaction with Y205^2.75^ that further linked the ECD and TMD. In addition to these polar interactions, there were extensive hydrophobic interactions between the receptor and each of the peptides; many of these interactions are conserved across other class B1 GPCRs, such as L141^1.36^, Y145^1.40^, Y148^1.43^, V194^2.64^, L201^2.71^, M233^3.36^, V237^3.40^, Y241^3.44^, W306^5.36^, R310^5.40^, I313^5.43^, L384^7.39^, E387^7.42^, L388^7.43^ (Figures 2, S4 and Table 2). Four of these conserved residues (Y145^1.40^, L201^2.71^, M233^3.36^ and L384^7.39^) were randomly selected for alanine mutagenesis and each individual mutant dramatically reduced the cAMP accumulation elicited by semaglutide or taspoglutide. With the exception of L384^7.39^A, these mutants had equivalent effects with GLP-1, suggesting that they play equivalent roles in different peptide-induced receptor activation (Figure S5).

As noted above, The C-terminus of the peptide was close to the ECD and ECL1 regions of GLP-1R, and the ECD-focused refinement enabled resolution to model the interactions between them (Figure S4). Generally, the side chain rotamer placements of the C-terminal halves of the peptides and their interactions with the receptor were similar for both structures (Figures 3, S4A and Table 2) and with our previously reported GLP-1 bound structure (Zhang et al. 2020). ECL1 had limited direct interactions with GLP-1 derived peptides, with the exception of W214^ECL1^ that exhibited π-π stacking with Trp31 in all three structures (Figures 3, S4A and Table 2) (Zhang et al. 2020). The ECD formed extensive hydrophobic interactions with the C-terminal halves of the peptides, including W39^ECD^, E68^ECD^, Y69^ECD^, Y88^ECD^, L89^ECD^, P90^ECD^ and W91^ECD^. In addition, hydrogen bonds were formed between Glu21 and Ser31^ECD^ at the N far terminus of the receptor, and between the backbone of Val33 and R121^ECD^, in all three structures (Zhang et al. 2020) (Figures 2, 3, S4A and Table 2). The importance of these residues in peptide binding and signalling of GLP-1R is supported by previous alanine mutagenesis studies (Wilmen et al. 1997, Day et al. 2011, Underwood et al. 2010). The C terminus of taspoglutide, beyond Arg34, extended the α-helical secondary structure of the peptide, however, for GLP-1 and semaglutide the helix terminated at residue 34 with the last two residues lacking secondary structure (Figure 3).

**Figure 3.**
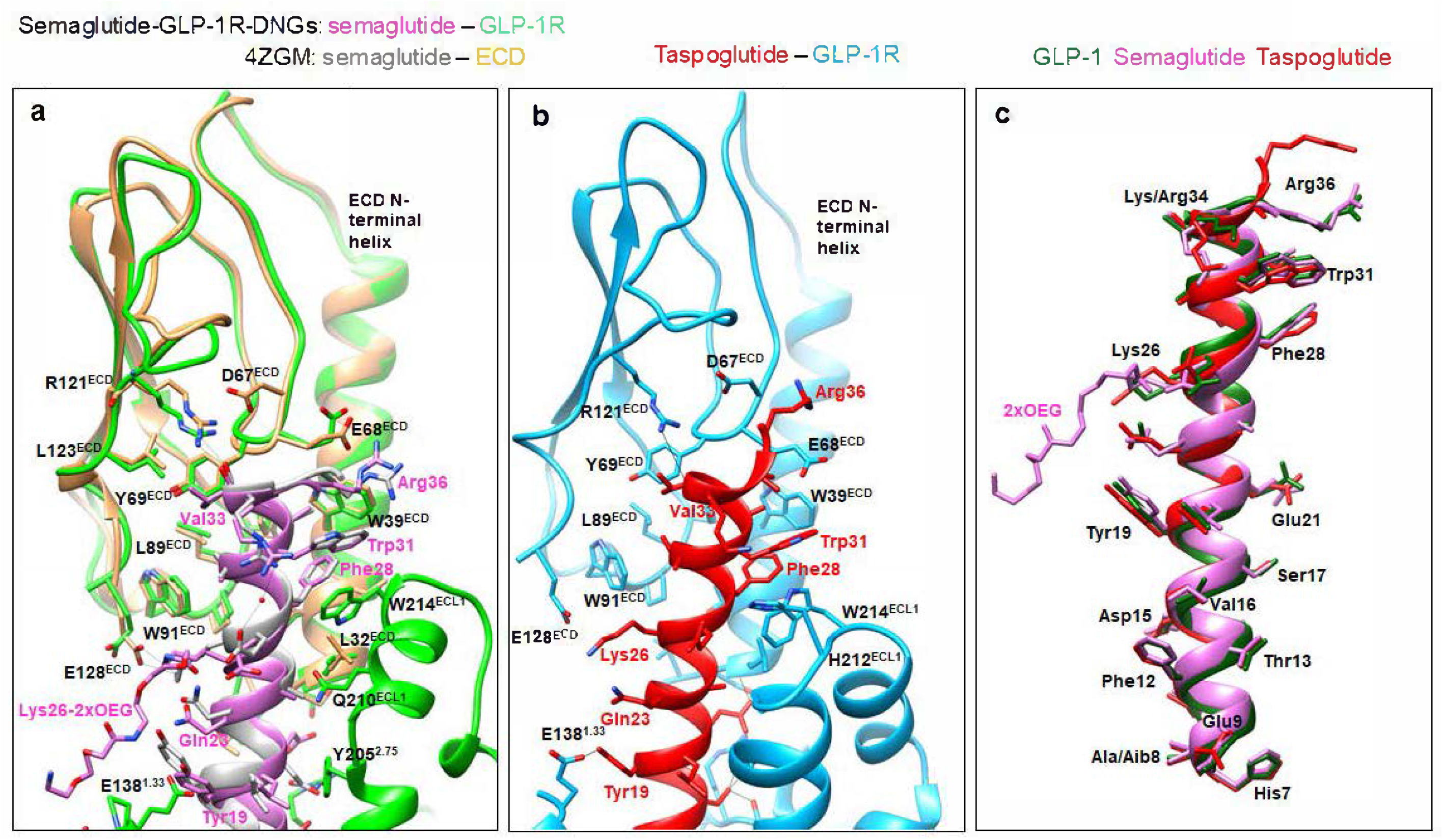
Interactions of semaglutide and taspoglutide with the GLP-1R ECD. GLP-1R residues that interact with peptide backbone or peptide side chains are displayed in stick format, coloured by heteroatom, with the backbone in ribbon format. **a**, Superimposition of the GLP-1R-Gs complex structure with the crystal structure of isolated semaglutide-GLP-1R ECD complex (PDB: 4ZGM); **b**, Taspoglutide-GLP-1R structure; **c**, Overlay of the GLP-1 (PDB: 6×18), semaglutide and taspoglutide peptide agonists. Colours are highlighted on the figure panels. Black lines depict hydrogen bonds as determined using UCSF chimera. *Also see Figures S4 and Table S2*.

In semaglutide, there is an Arg substitution of the Lys34 in GLP-1, however neither Arg34 of semaglutide or Lys34 of GLP-1 or taspoglutide formed interactions with the receptor (Zhang et al. 2020) (Figure S4A and Table 2), in line with previous reports of limited effect of Arg34 substitution on GLP-1R binding and signalling (Lau et al. 2015). Unlike GLP-1 and taspoglutide, semaglutide is acylated on Lys26 with a γGlu-2xOEG linker and C18 fatty diacid moiety (Figure S1A) that enables enhanced binding to plasma albumin and extended half-life in vivo (Lau et al. 2015). While this substitution was unresolved in the consensus map, density within the cryo-EM map was evident for the modified Lys26 within the ECD-focused map. Two distinct continuous densities could be observed that likely represent the most stable positions of the derivatised lysine (Figure S6A). Of the two densities, the one positioned down towards the top of TM1 had clearer density than the alternative density that extended up towards the receptor ECD. Nonetheless, only the linker Lys26-2xOEG moiety was able to be modelled in either orientation, before the density merged into the lower resolution regions of the detergent micelle or the receptor ECD (Figure S6B). Only the downward conformation of the Lys26-2xOEG moiety that had greater density was included in the final model. The structure of the C-terminal half of semaglutide and the receptor ECD closely resembles the crystal structure of semaglutide-bound GLP-1R ECD (PDB: 4ZGM) (Figure 3A). However, semaglutide Lys26 is not acylated in the ECD crystal structure (PDB: 4ZGM), and instead interacts with E128^ECD^ (Figure 3A); an equivalent interaction can be observed in GLP-1 and taspoglutide bound receptor structures that also have an unmodified Lys26 (Zhang et al. 2020) (Figure 3B). In the semaglutide-bound structure, the acylated Lys26 did not form this interaction (Figure 3). Semaglutide maintains high affinity and potency, which is consistent with the previous findings that substation of Lys26 or mutation of E128^ECD^ had limited effect on receptor affinity (Lau et al. 2015).

### Conformational comparison of different GLP-1 derived peptide-GLP-1R-Gs complexes

The GLP-1 and GLP-1 derived peptide complexes exhibited a similar global conformation for both receptor and Gs protein (Figure 4). The only notable distinction between taspoglutide- and GLP-1-bound GLP-1R in the consensus maps occurred within the top of TM2 and ECL1 region, whereas this region exhibited an identical conformation for semaglutide- and GLP-1 complexes (Figures 4A-B). Interestingly, the TM2/ECL1 conformation of the taspoglutide-GLP-1R structure is very similar to the previously published ExP5-GLP-1R structure (PDB: 6B3J), despite the distinct amino acid sequences between taspoglutide and ExP5 (Liang, Khoshouei, Glukhova, et al. 2018). Moreover, both peptides formed similar interactions with W214^ECL1^ and H212^ECL1^ of GLP-1R. This latter interaction was not observed in the GLP-1- and semaglutide-bound structures with the H212^ECL1^ sidechain located on the opposite side of ECL1 in these structures (Figure 4D).

**Figure 4.**
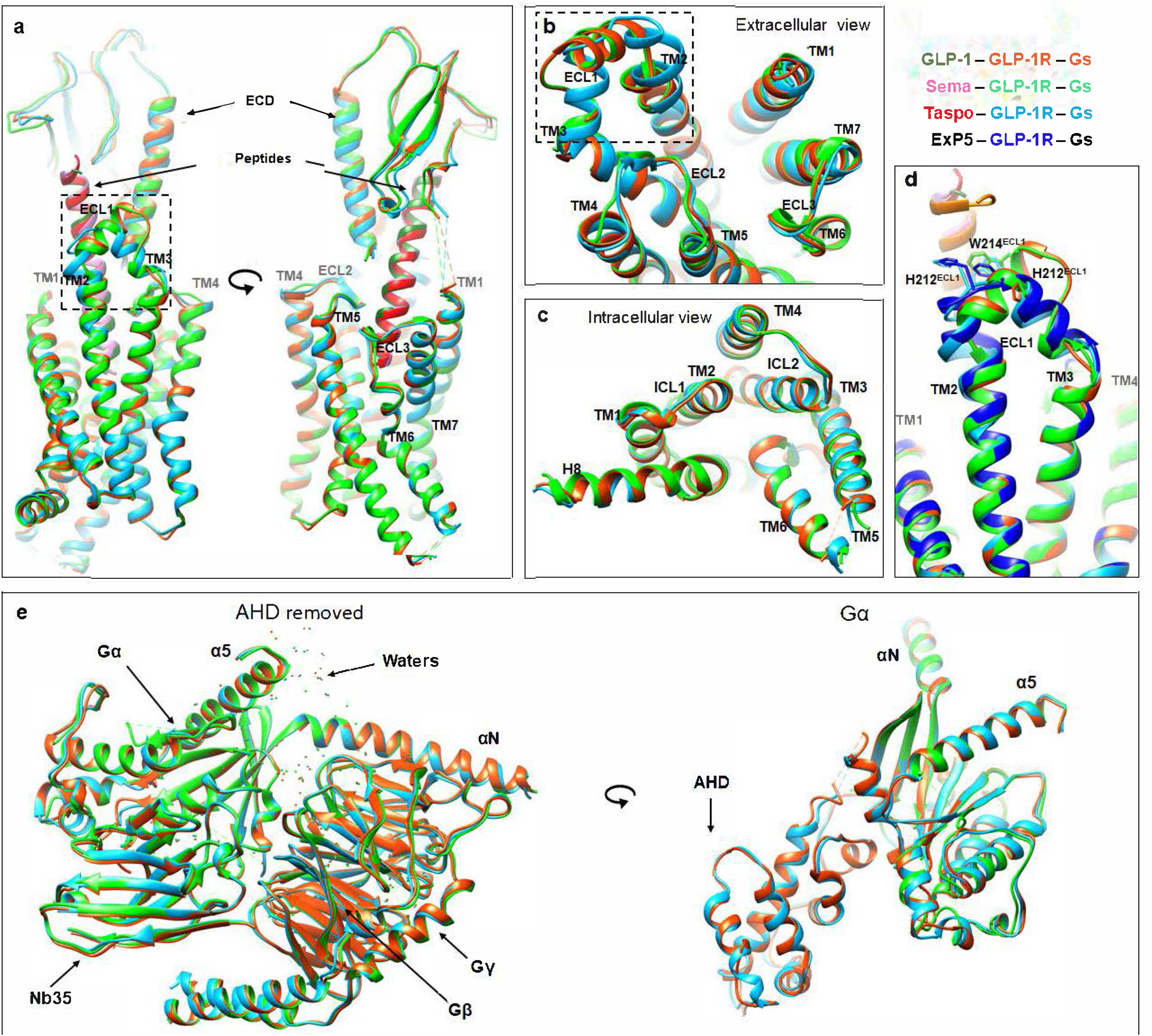
Peptide-bound GLP-1R-Gs complexes display similar active state conformations. **a-c**, Superimposition of the GLP-1 (PDB: 6×18), semaglutide and taspoglutide bound GLP-1R in backbone ribbon format from side view (**a**), extracellular view (**b**) and intracellular view (**c**); **d**, Superimposition of the GLP-1, semaglutide, taspoglutide and exendin P5 (ExP5; PDB: 6B3J) bound structures zoomed in to highlight the TM2/ECL1/TM3 region, with ECL1 residues that interact with bound peptides displayed in stick format; **e**, Superimposition of the G protein from GLP-1-, semaglutide- and taspoglutide-bound structures. Structures are displayed in backbone ribbon format coloured according to the receptor-ligand complex, with waters displayed as balls. Colours are highlighted on the figure panels. Regions of greatest divergence between the receptor structures are highlighted using dashed rectangles.

With the exception of TM2/ECL1, the metastable conformation of the receptor extracellular side, including the ECD and ECLs, was identical between the GLP-1 and GLP-1 derived peptides complexes (Figure 4) and all three receptors exhibited a similar reorganisation of the polar networks blow the peptide binding site and at the base of the receptor. These conformational changes facilitate the sharp kink at the centre of TM6 and a large outward movement of the intracellular half of TM6, characteristic of all the class B1 GPCR active state structures (Liang, Belousoff, Zhao, et al. 2020). Likewise, the rest of the intracellular side of the GLP-1R displayed an identical conformation (Figure 4C) and equivalent interactions with Gs, across the GLP-1 and GLP-1 derived peptide bound structures, including those interactions mediated by structural waters. Most of the observed interactions between the receptor and Gs protein were also consistent with other published GLP-1R-Gs structures, including those with small molecule agonists (Zhao et al. 2020, Liang, Khoshouei, Glukhova, et al. 2018, Zhang et al. 2020), which further supports a model of ligand-independent common interaction networks of the GLP-1R upon Gs protein coupling. The overall conformation of the G protein itself was also remarkably similar among the three structures (Figure 4E), including the structural waters within each subunit and their interfaces. Moreover, despite the high mobility of AHD of Gα subunit, the metastable conformation that was resolved in G protein-focused refinements in the taspoglutide-bound structure was similar to that previously described for the GLP-1 bound structure (Zhang et al. 2020) (Figure 4E).

### 3D variability analysis reveals distinct dynamic motions in different GLP-1 derived peptide complexes

As described above, the static consensus structures for taspoglutide, semaglutide and GLP-1 bound, active GLP-1Rs are remarkably similar. For other class B1 GPCRs, we have demonstrated that the active complexes are dynamic and that dynamics can contribute to the pharmacological behaviour of bound ligands (Dong et al. 2020, Liang, Belousoff, Fletcher, et al. 2020, Zhang et al. 2020). To understand and visualize the dynamic motions in GLP-1R complexes, 3D variability analysis implemented in cryoSPARC was performed. In 3D variability analysis, particles of individual complex were partitioned into the top five principal components (modes) and reconstructed into a continuous series of 20 3D volumes (frame000 – frame019), each mode corresponding to a different type of variability. The top three principal components of both semaglutide- and taspoglutide-bound GLP-1R-Gs complexes were recorded in Video S1, and revealed common twisting and rocking motions of the GLP-1R relative to the Gs protein (Video S1 component 2 and 3), where the extent of motion was comparable to that previously reported for GLP-1-bound complex (Zhang et al. 2020). However, clear differences in the dynamics of each complex were observed, with the most notable changes occurring at the extracellular side of the receptor, as shown in Video S1 component 1. The extracellular end of TM2 and ECL1 of the semaglutide-bound receptor exhibited a robust motion, consistent with the lower resolution of this region compared to maps with the other peptide (Figures S1B-C). Modelling the backbone of the GLP-1R into the two maps at the extremes of the mode (frame000 and frame019) of component 1 revealed a 7 Å and 14 Å movement of the top of TM2 and ECL1, respectively (Figure 5A). This region was also dynamic in GLP-1-bound GLP-1R, but with less movement (limited movement of the top of TM2 and only a 7 Å movement of ECL1) (Zhang et al. 2020). Moreover, differences were observed in the direction of motion between the GLP-1- and semaglutide-bound structures. In the GLP-1-bound GLP-1R, ECL1 moved towards TM1 (Zhang et al. 2020), whereas there was movement towards TM1 linked with a large outward movement in the semaglutide-bound structure (Video S1, Figure 5A). In contrast, the TM2/ECL1 region of taspoglutide-bound GLP-1R was relatively stable compared to that of the semaglutide and GLP-1 complexes (Video S1, Figure 5B). Interestingly, while the extracellular portions of TMs1/6/7 and ECL3 were relatively stable in the complex with GLP-1 (Zhang et al. 2020), these regions underwent a large motion away from the TM core in the semaglutide complex, transitioning 7 Å and 8 Å at the top of TM1 and TM7, respectively (Video S1, Figure 5A). Moreover, the extracellular tip of TM4/5 and ECL2 also moved outward. In parallel, the whole semaglutide peptide coordinately moved almost 5 Å towards TMs1/6/7, with the exception of the very end of its N terminus (Video S1, Figure 5A). A similar, but less dramatic, motion was observed in the taspoglutide-bound structure, with 3 Å movement of the taspoglutide peptide towards TMs1/6/7, facilitated by a smaller outward motion of TM1 top, TM6/ECL3/TM7 and TM4/ECL2/TM5 relative to the remainder of the TM bundle (Video S1, Figure 5B).

**Figure 5.**
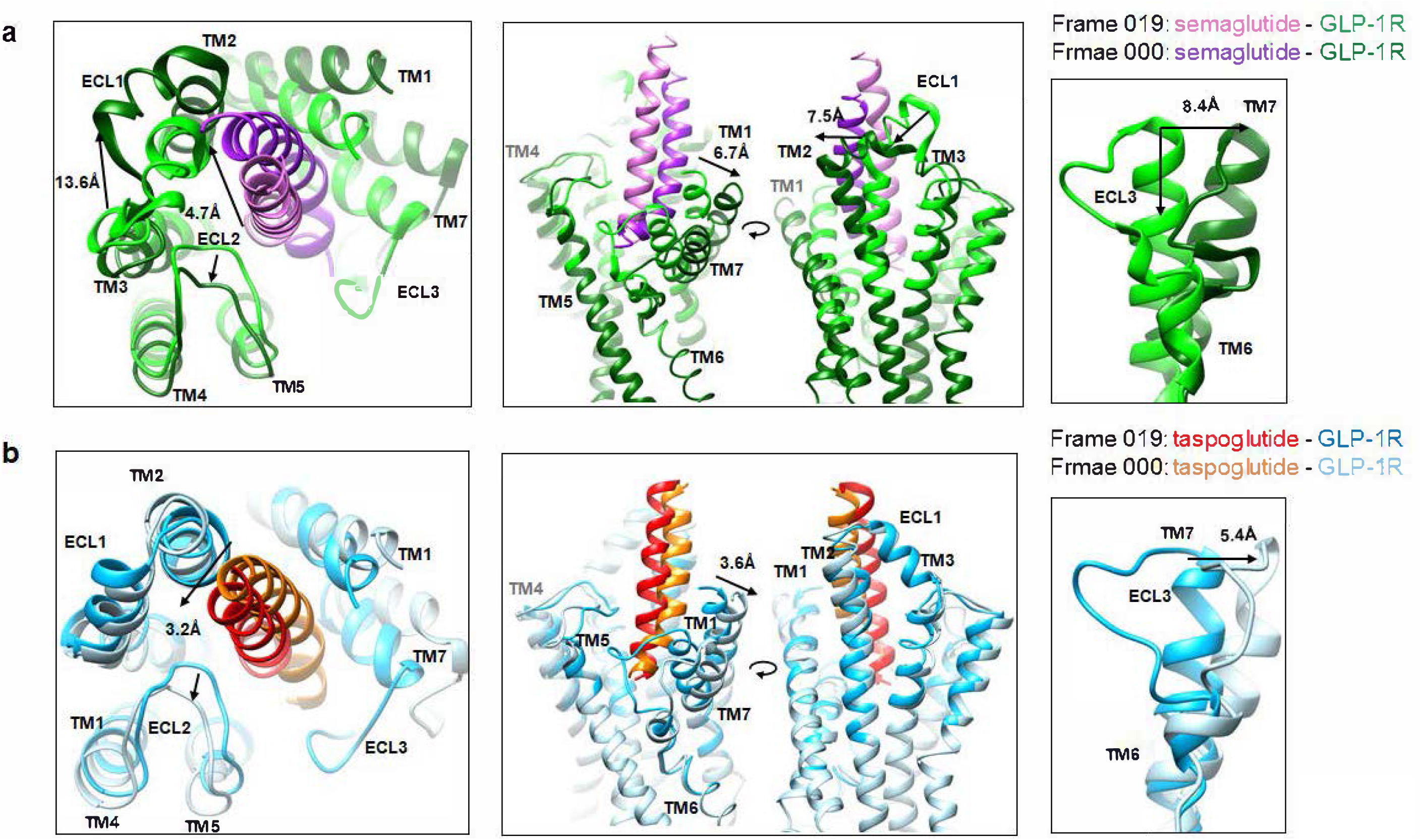
GLP-1R conformational transitions observed in 3D variability analyses of the semaglutide-(a) or taspoglutide-(b) bound structures. Static backbone models were built into the cryo-EM maps from frame 000 and frame 019 of component 1 in Video S1 and superimposition of these two extreme models of the complex displayed from top view (Left), side view (Middle) and zoom in of the TM6/ECL3/TM7 region (Right). **a**, Semaglutide-GLP-1R-Gs complex. The distance between the top of TMs1/7 in the two extreme maps was 6.7 Å and 8.4 Å when measured at Cα of E139^1.34^ and G377^7.32^, respectively. Motions of TM2 and EC1 were observed spanning 7.5 Å and 13.6 Å when measured at Cα of M204^2.74^ and D215^ECL1^, respectively. The peptide underwent a 4.7 Å movement measured between Cα of Val33 (frame 000) and Ala30 (frame 019). **b**, Taspoglutide-GLP-1R-Gs complex. The motions of top of TMs1/7 were 3.6 Å and 5.4 Å when measured at Cα of E139^1.34^ and G377^7.32^, respectively. The peptide had a 3.2 Å movement measured between Cα of Ala30 (frame 000) and Ile29 (frame 019). Colours are highlighted on the figure panels. *Also see Figure S7 and Videos S1 and S2*.

The extent of motion of the extracellular side of TM bundle and peptide N-terminus corresponded to the relative change in the ECD (Video S2 component 4), consistent with the lower resolution of the ECD relative to the remainder of the receptor in both semaglutide- and taspoglutide-bound structures (Figure S1B-C). The ECD of the taspoglutide-bound receptor shifted 12 Å when measured between the Cα of G52^ECD^ in the models built in the maps at the extremes of component 4 (Figure 6A), compared to a 7 Å motion of the ECD in GLP-1-bound structures through an equivalent analysis (Zhang et al. 2020). The ECD of the semaglutide-bound GLP-1R was more dynamic than the other peptides (Videos S1 & S2), leading to ambiguous backbone modelling in the extreme map of the dynamic state (frame000), and thus the ECD was rigid body fitted into frame000 map of component 4 (Figures 6A, S7B). As the metastable position of the ECD, particularly the N terminal helix, was identical among these structures (Figure 4A), the consensus semaglutide-bound GLP-1R model was shown as a reference to compare the direction of motion of the ECD in the receptor complexes with different peptides. In the semaglutide-bound structure, the ECD shifted between the consensus position and TM1, while it moved towards ECL1 in the GLP-1- and taspoglutide-bound structures, with the angle between the two transitional routes around 90° (Figure 6B).

**Figure 6.**
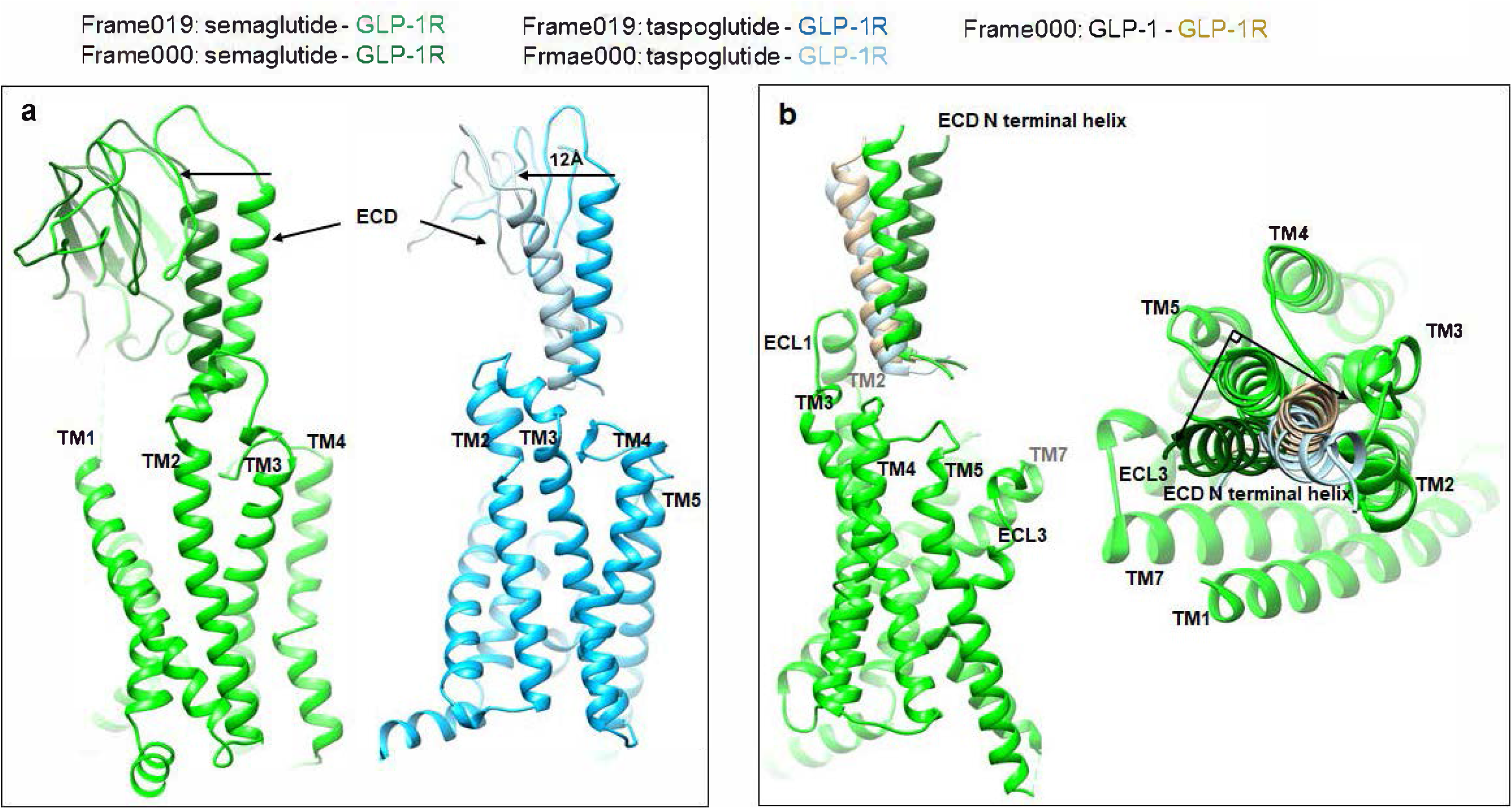
GLP-1R ECD conformational transitions observed in 3D variability analyses of the semaglutide- or taspoglutide-bound structures. Static backbone models, displayed in ribbon format, were built into the cryo-EM maps from frame 000 and frame 019 of component 4 in Video S2. **a**, Dynamic analysis revealed a 12 Å transition between the maps (when measured from the Cα of G52^ECD^) for taspoglutide-bound complexes. **b**, Superimposition of the consensus structure of semaglutide-bound GLP-1R (frame 019) and dynamic states (frame of semaglutide-, taspoglutide- and GLP-1-bound ECD N terminal α-helix structures. A 93° angle difference was observed between Cα of G52^ECD^ in consensus structure, and the location of this residue in the dynamic state of semaglutide-versus GLP-1-bound receptor ECD. Colours are highlighted on the figure panels. *Also see Figure S7 and Video S1*.

The G protein heterotrimer (with the exception of AHD) in the presence of Nb35 was relatively stable with only slight motions at the protein surface (Video S1). In contrast, the AHD was highly mobile, shifting between an open position and a more closed positon relative to the ras-like domain of Gαs subunit in the taspoglutide-bound complex (Video S2), similar to previous observations with GLP-1-bound structure(Zhang et al. 2020). However, in the semaglutide complex, the AHD adopted only a single open conformation in the first three principle components, while in the less populated components (component 4 and 5), a weak density of a more closed form could be observed (Video S2). Concurrently, 3D classification in RELION was performed to determine the most abundant, stable, AHD positions and the relative population of structures in different positions. Similar to the GLP-1-bound complex, the AHD in the taspoglutide-bound complex displayed three major classes exhibiting an open, middle and closed conformation (Figure 7B). Although there were more particles for the AHD in the open conformation (51%), the closed form containing only 14% of the particles was the best resolved enabling this to be improved in the G protein-focused refinement and, subsequently, for this region to be modelled at a backbone level (Figure S1C). The distribution of particles in the five classes of the semaglutide complex, however, was very different to that of the taspoglutide-bound complex, with two classes (19% and 28% particles) occupying the open position, and one class of 17% particles in a location similar to the middle position of AHD in the taspoglutide-bound complex. The other two classes were largely varied and appeared to cross positions between different classes (Figure 7A). Moreover, none of these classes was well resolved. A morph between the major conformations was recorded in Video S2.

**Figure 7.**
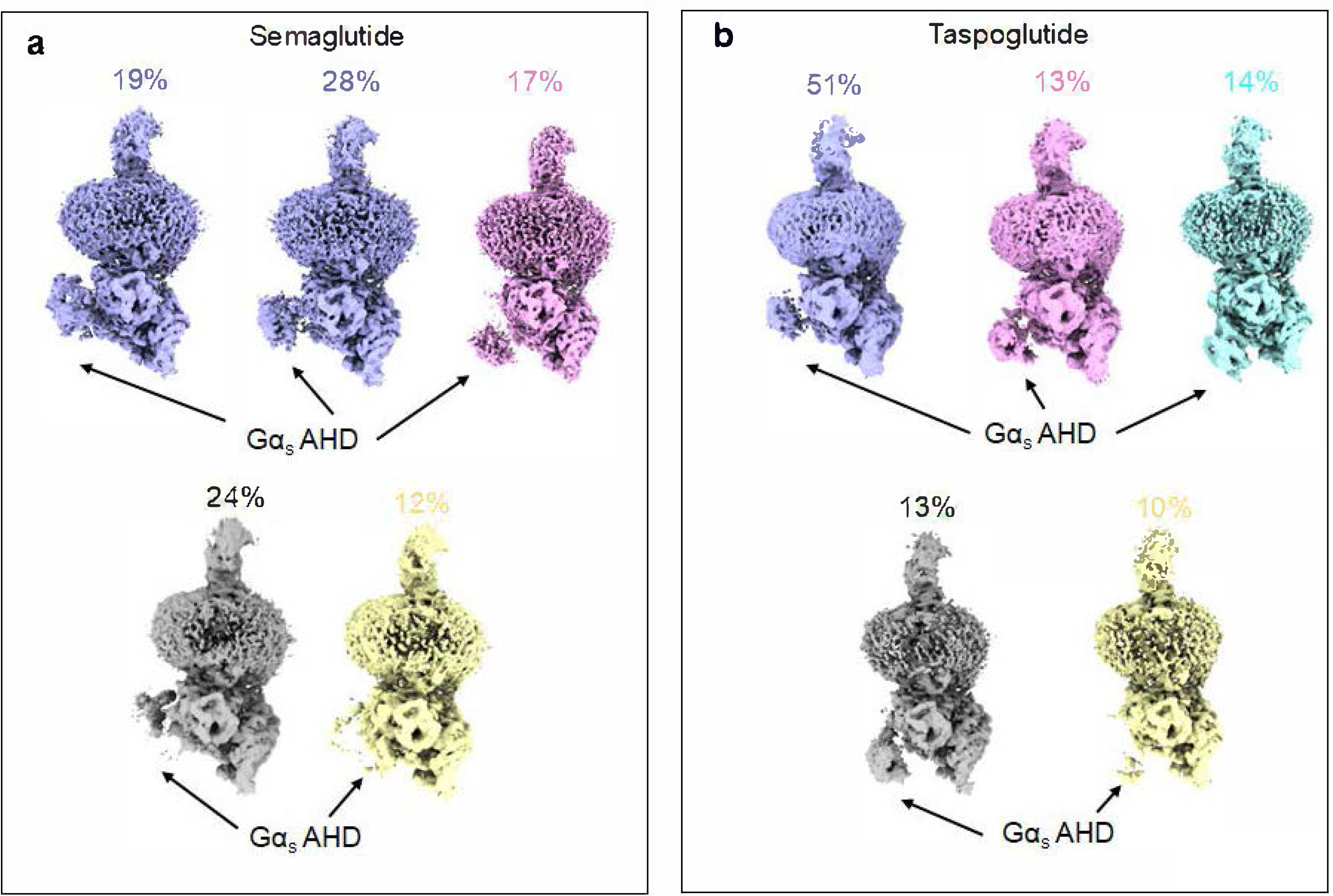
AHD conformational differences identified by focused 3D classification in RELION. Multiple conformations from 3D classification in RELION were identified, with three major orientations, “open” (purple), “intermediate” (pink), and “closed” (cyan) observed. The other two classes differed between complexes and had less well-defined AHDs (grey and yellow). **a**, Semaglutide-bound complex; Two classes have an “open” conformation, and one with an “intermediate” conformation. **b**, Taspoglutide bound complex; The AHD adopted three major conformations. The percentage of particles contributing to each class is shown above the density maps, labelled in the same colour. *Also see Video S2*.

## DISCUSSION

Taspoglutide has 93% identity with endogenous GLP-1 (7-36)NH_2_, with replacement of amino acids at position 8 and 35 by Aib to improve the DDP-4 and serine protease resistance; the latter substitution was also designed to increase the C-terminal helicity to increase the receptor binding affinity (Sebokova et al. 2010). In line with this, continual helical secondary structure for the far C-terminus of taspoglutide could be resolved, whereas the C terminus (Gly35-Arg36) of GLP-1 and semaglutide lack secondary structure (Figure 3C). Despite a comparable clinical efficacy for glycaemic control and weight loss to approved GLP-1R agonists, the development of taspoglutide was terminated in late phase 3 clinical trials due to severe gastrointestinal side effects and systemic allergic reactions (Rosenstock et al. 2013). In contrast, once-weekly semaglutide, approved in 2017, was designed to maintain structural similarity to endogenous GLP-1 but with reduced DPP4 metabolism and enhanced plasma albumin binding to decrease clearance (Lau et al. 2015). This is achieved by Aib8 substitution, and replacement of Lys34 with Arg to enable site-specific acylation of Lys26 (Lau et al. 2015). Comparison of the GLP-1- and semaglutide-bound GLP-1R structures revealed that the Arg34 substitution did not modify the secondary structure of the peptide C terminus (Figure 3C). While density for the modified Lys26 was not resolved in initial maps, the linker could be resolved by focused refinement of the ECD. The most prominent density for the linker was oriented towards the detergent micelle, suggesting that the lipid component extends into the micelle and likely interacts with the lipid bilayer in native membranes, stabilising the downward position of the lysine and linker.

Despite these minor differences in peptide structure, semaglutide and taspoglutide share high sequence identity with GLP-1, and they exhibited similar potency in cAMP accumulation compared to GLP-1 in CHO cells that overexpress (Figure S5). Thus, it is not surprising that the peptide binding mode, and global conformations of the receptor and Gs protein were very similar for their active structures (Figures 2-4). Indeed, in the consensus models, the only notable difference in receptor structure occurred in ECL1 (S206-S219) that adopted different conformations between taspoglutide- and semaglutide-bound complexes, with the latter similar to that of the GLP-1-bound structure (Zhang et al. 2020). Nonetheless, only limited direct interactions occurred between ECL1 and the peptides (Figure 3 and Table S2). This is consistent with earlier alanine mutagenesis studies, where individual mutation of S206-S219 had only limited effect on GLP-1-induced receptor affinity and cAMP signalling (Wootten, Reynolds, Smith, et al. 2016, Yang et al. 2016), albeit that further structure-function studies of this region for semaglutide, and in particular, taspoglutide, are required to understand the importance of this region for their function.

Semaglutide has become the gold standard for GLP-1R agonists demonstrating more robust effects in control of glycaemia and body weight than other GLP-1R agonists, with a comparable safety profile (Tsoukas et al. 2017, Ahmann et al. 2018, O’Neil et al. 2018, Pratley et al. 2018). Moreover, semaglutide has greater efficacy in reducing cardiovascular risk than the other currently marketed peptides (Marso et al. 2016) and holds promise for the treatment of non-alcoholic steatohepatitis (NASH) where it is in phase 2 clinical trials (NCT02970942). In contrast taspoglutide did not reduce cardiovascular risk (Henry et al. 2012). While only limited pharmacological data is currently available for taspoglutide, both semaglutide and taspoglutide are reported to have distinct recycling and trafficking profiles relative to GLP-1 (Fletcher et al., 2019). Consequently, there must be distinct mechanisms that contribute to these pharmacological differences beyond the metastable consensus structures that exhibit almost identical patterns of interactions between the peptides and the receptor. As accumulated data suggests the GPCR dynamics may influence their functions (Dong et al. 2020, Liang, Belousoff, Fletcher, et al. 2020, Zhang et al. 2020), we further analysed the conformational dynamics of semaglutide- or taspoglutide-bound active complexes.

The ability of cryo-EM to capture spectrums of conformations present during vitrification allows insight into conformational dynamics of proteins (Lau et al. 2018, Murata and Wolf 2018), and was critical in the identification of distinctions in the behaviour of the different GLP-1R peptide complexes. The potential for differences in the mobility of the complexes could be inferred from the relatively lower resolution for the ECD, extracellular portion of the TM domain and the AHD of Gαs, suggesting that these domains were more dynamic. Therefore, we hypothesized that the conformational dynamics of GLP-1R complexes, rather than the consensus metastable interactions, could contribute to their distinct pharmacological profiles.

Parsing out the principal components contributing to different conformational ensembles using 3D variance analysis revealed that the extracellular side of TMs1/6/7 and ECL3 underwent a large movement away from the TM core in semaglutide structure (Figure 5A), while this region was relatively stable in the GLP-1-bound structure (Zhang et al. 2020). Active GLP-1R structures in complex with the biased agonists, CHU-128, TT-OAD2 and ExP5 have different conformations of this region relative to that of GLP-1-bound GLP-1R (Zhao et al. 2020, Liang, Khoshouei, Glukhova, et al. 2018, Zhang et al. 2020), and previous mutagenesis studies have shown that this region is crucial for biased agonism (Wootten, Reynolds, Smith, et al. 2016, Lei et al. 2018). In parallel with the conformational change in TM1/6/7/ECL3, there was outward motion of TM2/ECL1/TM3 and TM4/ECL2/TM5, leading to enlargement of the binding cavity and a 5 Å shift of the semaglutide peptide towards TMs1/6/7, resulting in the loss of interactions with ECL2 (Figure 5A). Extensive mutagenesis studies of the GLP-1R have suggested the ECL2 interactions are essential for all peptide-induced receptor activation (Koole et al. 2012, Dods and Donnelly 2015, Wootten, Reynolds, Smith, et al. 2016, Yang et al. 2016). While the outward motions were also observed in the taspoglutide-bound complex, the extent of motion was lower. We speculate that the “open” state might represent an intermediate conformation that occurs during peptide-receptor association / dissociation.

The ECD in the taspoglutide-bound structure was also less dynamic than that of the semaglutide-bound receptor, and this was linked to stronger peptide interactions with ECL1 that limit the motion of the peptide, as well as the dynamics of ECL1 (Videos S1 & S2). The ECD in both structures underwent a twisting motion that is consistent with the predicted motion of the ECD as it moves from an inactive state to agonist-bound states (Zhang et al. 2020), however, this motion was greater in the semaglutide-bound complex, leading to a shift in the position of the N terminal helix of the ECD from the consensus position towards TM1 (Figure 6B). In contrast, the more restricted ECD motion in the taspoglutide-bound GLP-1R moved the N-terminal helix towards ECL1. The direction of motion for the taspoglutide complex was more consistent with that seen for the GLP-1-bound GLP-1R ECD, and also that of TT-OAD2 (Zhang et al. 2020); ∼90° different from the semaglutide complex (Figure 6B). Previous work has established a critical role for the GLP-1R ECD in peptide-induced cAMP signalling (Yin et al. 2016), however, how the ECD dynamics influence the cell signalling is still unknown. Nonetheless, recent work on CGRP/adrenomedullin receptors has provided evidence that the ECD dynamics of these receptors plays a key role in peptide selectivity (Liang, Belousoff, Fletcher, et al. 2020). Intriguingly, although equivalent 3D variance analysis was not performed, multiple 3D classes that differed in the location of the ECD were resolved for the active complex of the human parathyroid hormone receptor-1 (Zhao et al. 2019). In this study, the data were suggestive of separation of peptide C-terminus and the ECD in one of the classes, but with the peptide N-terminus still engaged with the TMD core. Whether an equivalent process also occurs for the GLP-1R is unclear, but the loss of resolution where larger motions were observed could be consistent with partial disengagement, and the dynamics of the complexes are likely to play an important role in peptide dissociation.

The AHD of the Gα subunit has been poorly resolved in most cryoEM structures of GPCRs (Maeda et al. 2019, Draper-Joyce et al. 2018, Qiao et al. 2020, Liang, Belousoff, Zhao, et al. 2020, Liang, Belousoff, Fletcher, et al. 2020, Zhao et al. 2019, Liang, Khoshouei, Deganutti, et al. 2018, Liang et al. 2017), indicative of high mobility of this region in the nucleotide-free state and conditions of complex formation. Multiple conformations of AHD were observed in low resolution 3D classes of active β2-adrenoceptor and calcitonin receptor Gs complexes (Liang et al. 2017, Westfield et al. 2011). Our recently published high-resolution structures of GLP-1R-Gs complex with GLP-1, PF 06882961 or CHU-128 revealed open, intermediate and closed conformations of AHD relative to the Gα ras-like domain following focused 3D classification, and the AHD in the closed position was best resolved, albeit not the most abundant, for all three structures (Zhang et al. 2020). A similar distribution of metastable AHD positions was observed in the taspoglutide-bound complex in the current study (Figure 7B). In contrast, equivalent analysis of the semaglutide-bound complex suggested a more varied (dynamic) AHD and no closed conformation was observed (Figure 7A). Nonetheless, the 3D variability analysis provided confirmatory data that the AHD in the semaglutide-bound complex was predominantly present in the open conformation, albeit that weak density for other conformations could be visualised, and this contrasted to the continuous motion from open to closed conformations that could be observed in the taspoglutide-bound GLP-1R (Video S2). While there are caveats on the extrapolation of the structural data, including the modifications to Gs, presence of Nb35 and detergent micelle environment, biophysical studies have previously demonstrated ligand-dependent differences in conformational sampling of Gs that could be linked to GTP binding and G protein turnover (Furness et al. 2016). As such, the differences in observed conformational dynamics of Gs in our structures could provide insight into efficacy differences between ligands.

In conclusion, we have solved 2.5 Å cryoEM structures of GLP-1R-Gs complexes with two peptides, taspoglutide and semaglutide, that had distinct outcomes in clinical trials. These revealed highly similar metastable peptide binding modes and conformations of the active receptor-Gs complex that were also similar to that of the native GLP-1 peptide. However, each GLP-1R complex displayed unique peptide-dependant dynamics suggesting that conformational dynamics may be critical for pharmacological differences between the peptides. While much more structural data, including complexes with different transducers, and complexes in lipidic environments, will be required to fully understand the link between structure and function, our structures and the dynamic information captured by cryo-EM provide mechanistic insights into the ligand binding, receptor activation and G protein coupling of class B1 GPCRs.

## ACKNOWLEDGEMENTS

This work was supported by the National Health and Medical Research Council of Australia (NHMRC) (project grant 1126857, ideas grant 1184726 and SRF 1160076 (D.W.)), and program grant 1150083 (P.M.S.)). P.M.S. is a Senior Principal Research Fellow and D.W. a Senior Research Fellow of the NHMRC. R.D. was supported by Takeda Science Foundation 2019 Medical Research Grant and Japan Science and Technology Agency PRESTO (18069571). This work was supported by the Monash University MASSIVE high-performance computing facility.

## AUTHOR CONTRIBUTIONS

**Conceptualization**: D.W. and P.M.S. conceived the project and formulated the research plan. **Investigation**: X.Z. expressed and purified receptor complexes, generated atomic models and performed the pharmacological assays; R.D performed sample vitrification and cryo-EM imaging; X.Z and R.D processed the EM data. **Formal Analysis**: X.Z., D.W., P.M.S., M.B. analysed the receptor structures; D.W and X.Z. analysed pharmacological data. **Supervision**: D.W. and P.M.S supervised the whole project; Y-L.L assisted with supervision for complex biochemistry; M.B supervised X.Z in cryo-EM processing and crosschecked the molecular models. **Project administration**: D.W, P.M.S and M.B coordinated the biochemistry, pharmacological studies, data processing and analysis; R.D coordinated sample vitrification and cryo-EM imaging. **Visualization**: X.Z prepared figures and videos for the manuscript. **Writing - original draft:** X.Z, P.M.S and D.W wrote/contributed to the original manuscript draft. **Writing - review & editing:** M.B and R.D contributed to writing, review and editing of the manuscript. **Funding Acquisition**: D.W., P.M.S. and R.D. acquired the financial support for the project leading to this publication.

## STAR methods

### CONTACT FOR REAGENT AND RESOURCE SHARING

Further information and requests for resources and reagents should be directed to and will be fulfilled by the Lead Contact, Denise Wootten.

### EXPERIMENTAL MODELS

Protein expression for biochemical analysis and purification was performed in *Spodoptera frugiperda (Sf9)* and *Trichuplusia ni* (*Tni*) insect cells, maintained in ESF 921 serum free media (Expression systems) at 30°C. CHOFlpIn cells used for mammalian cell studies were maintained in DMEM supplemented with 5% FBS at 37°C in 5% CO_2._ *Escherichia Coli* (*E. coli*) strain BL21, used to express Nb35, was cultured in Luria-Bertani (LB) liquid medium (10 g tryptone, 10 g NaCL and 5 g yeast extract per litre) with continuous shaking (180 rpm) or on LB agar plate (LB medium with 15 g agar per litre) at 37°C.

## METHOD DETAILS

### Constructs

The human GLP-1R was modified to replace the native signal peptide with that of haemagglutinin (HA) to improve receptor expression and contain N-terminal Flag tag epitope and C-terminal 8xHis tag. 3C protease cleavage site (LEVLFQGP) was inserted between both tags and the receptor. The construct were generated in both insect and mammalian cell expression vectors, and the modification did not affect the receptor pharmacological profile (Liang, Khoshouei, Glukhova, et al. 2018). Dominant negative Gαs (DNGαs) includes mutations S54N, G226A, E268A, N271K, K274D, R280K, T284D, I285T and A366S of Gαs, which has been described preciously to reduce nucleotide binding affinities and enhance the stability of GLP-1R and Gs heterotrimer complex(Liang, Belousoff, Fletcher, et al. 2020, Liang, Zhao, et al. 2018). 8xHis tagged Gβ1-γ2 construct and 8xHis tagged Nb35 were provided by B.Kobilka(Rasmussen et al. 2011).

### Insect cell expression

Human GLP1R, human DNGαs and human Gβ1-γ2 constructs were co-expressed in Tni insect cells using baculovirus expression system as previously described(Liang, Khoshouei, Glukhova, et al. 2018). Briefly, Tni insect cells were grown in ESF 921 serum-free media to a density of 3.5 million/mL and infected with GLP1R, DNGαs and Gβ1-γ2 baculovirus at multiplicity of infection (MOI) ratio of 3:3:1. Culture was harvested by centrifugation 48 h post infection and cell pellets were stored at −80°C.

### Complex formation and purification

GLP-1R-DNGs complex formation and purification were performed as described by Liang, *et al*.(Liang, Khoshouei, Glukhova, et al. 2018) Cell pellets (from 1L insect cell culture) were thawed and suspended in 20mM HEPES pH 7.4, 50mM NaCl, 5mM CaCl_2_, 2mM MgCl_2_ supplemented with cOmplete Protease Inhibitor Cocktail tablets (Sigma Aldrich) and benzonase (Merk Millipore). Complex was formed by adding GLP-1R agonists (10µM semaglutide or 10µM taspoglutide, China Peptides), Nb35–His (10µg/ml) and apyrase (25mU/ml, NEB); the suspension was incubated for 1h at room temperature. The complex was solubilized from membrane using 0.5% (w/v) lauryl maltose neopentyl glycol (LMNG) and 0.03% (w/v) cholesteryl hemisuccinate tris salt (CHS) (Anatrace) for 1h at 4 °C. Insoluble material was removed by centrifugation at 30,000g for 20 min and the solubilized complex was immobilized by batch binding to M1 anti-Flag affinity resin in the presence of 5mM CaCl_2_.

The resin was packed into a glass column and washed with 20 column volumes of 20 mM HEPES pH 7.4, 100mM NaCl, 2mM MgCl_2_, 5mM CaCl_2_, 1µM peptide, 0.01% (w/v) LMNG and 0.0006% (w/v) CHS followed by elution with buffer containing 5mM EGTA and 0.1mg/ml FLAG peptide. The complex was then concentrated using an Amicon Ultra Centrifugal Filter (MWCO, 100 kDa) and subjected to size-exclusion chromatography (SEC) on a Superdex 200 Increase 10/300 column (GE Healthcare) that was pre-equilibrated with 20mM HEPES pH 7.4, 100mM NaCl, 2mM MgCl_2_, 1µM peptide, 0.01% (w/v) LMNG and 0.0006% (w/v) CHS to separate complex from contaminants. Eluted fractions consisting of receptor and G-protein complex were pooled and concentrated to 2-4mg/mL. The complex samples were flash frozen in liquid nitrogen and stored at −80°C.

### SDS-PAGE and Western blot analysis

Samples from important steps during purification were collected and analysed by SDS–PAGE and western blot. TGX™ Precast Gel (BioRad) was used to separate proteins within samples at 200V for 30min. Then gels were either stained by Instant Blue (Sigma Aldrich) or immediately transferred to PVDF membrane (BioRad) at 100V for 1h. The proteins on the PVDF membrane were probed with two primary antibodies simultaneously, rabbit anti-Gs C-18 antibody (cat. no. sc-383, Santa Cruz) against Gαs subunit and mouse poly-His antibody (cat. no. 34660, QIAGEN) against His tag. The membrane was washed and incubated with secondary antibodies followed by probing with FLAG-FITC antibody (prepared in the lab) against Flag tag on GLP-1R. The membranes were imaged using Typhoon 5 imaging system (Amersham).

### Negative staining and data processing

The complex samples were diluted to 0.006mg/mL in 20mM HEPES pH 7.4, 100mM NaCl, 2mM MgCl_2_ and 1µM peptide and applied to the continuous carbon grids (EMS). The grids were stained with 0.8% (w/v) uranyl formate solution and imaged using cryo-EM Tecnai™ T12 TEM at 120 kV. Around 50 images for each complex were collected with a magnified pixel size of 2.06 Å, and ∼20,000 particles of each complex were auto picked, extracted and 2D classified by RELION-3.0-beta.

### Vitrified sample preparation and data collection

Samples (3µL) were applied to glow-discharged Quantifoil R1.2/1.3 Cu/Rh 200 mesh grids (Quantifoil GmbH, Großlöbichau, Germany) and were flash frozen in liquid ethane using a Vitrobot mark IV (Thermo Fisher Scientific, Waltham, Massachusetts, USA) set at 100% humidity and 4°C for the prep chamber and 10 s blot time. Data were collected on a Titan Krios G3i microscope (Thermo Fisher Scientific) operated at an accelerating voltage of 300 kV with a 50 μm C2 aperture at an indicated magnification of 105K in nanoprobe EFTEM mode. Gatan K3 direct electron detector positioned post a Gatan BioQuantum energy filter (Gatan, Pleasanton, California, USA), operated in a zero-loss mode with a slit width of 25 eV was used to acquire dose fractionated images of the samples with a 100 µm objective aperture. Movies were recorded in hardware-binned mode (previously called counted mode on the K2 camera) with the experimental parameters listed in Table S1 using a 9-position beam-image shift acquisition pattern by custom scripts in SerialEM(Schorb et al. 2019).

### Data processing

As Figure S2 shows, 7488 and 8759 movies of semaglutide- and taspoglutide-GLP-1R-DNGs complexes, respectively, were motion corrected using MotionCor2(Zheng et al. 2017) and subjected to contrast transfer function (CTF) estimation using Kai Zhang’s GCTF software(Zhang 2016). Particles were picked from corrected micrographs using cryOLO software(Wagner et al. 2019) for the semaglutide dataset and RELION (version 3.0.7)(Zivanov et al. 2018) reference-based picker with a 20 Å low-pass filtered 3D map reference for the taspoglutide dataset. Picked particles were extracted and 2D classified using RELION (version 3.0.7). The selected 2D particles were used to generate the initial 3D model based on Stochastic Gradient Descent (SGD) algorithm, and subsequently applied to 3D classification. Particles from the best-looking class were subjected to Bayesian particle polishing, 2D classification, CTF refinement and cycles of 3D auto-refinement in RELION. 886,738 and 625,241 particles for semaglutide- and taspoglutide-GLP-1R complex were chosen to generate a final map using 3D auto-refinement and sharpened with a B-factor of −55Å and −70Å, respectively. Local resolution was determined using RELION with half-reconstructions as input maps. The map density of detergent micelle and Gαs helical domain was averaged out during the final map reconstruction for clarity.

To improve the local map quality, each refined particle was subjected into another 3D classification with a loose mask of ECD in RELION. The best resolved classes, 233K and 346K particles of semaglutide and taspoglutide complexes respectively, were further refined to generate ECD-focused maps. The refined particle stack was also 3D classified into 5 classes using a broad mask of the AHD without alignment. Each particle class (244K, 204K, 168K, 144K or 104K particles of semaglutide complex; 317K, 85K, 84K, 79K or 61K particles of taspoglutide complex) was 3D auto-refined, and the best resolved classes of taspoglutide complex (158K particles) was further refined and post processed with a G protein mask to generate a final map of the AHD.

### Atomic model refinement

The model of GLP-1-GLP-1R-DNGs (PDB:6×18) used as initial template was fitted in the cryo-EM density maps with the MDFF routine in namd2(Chan et al. 2012). The fitted model for the GLP-1R transmembrane domain (TMD), Gs protein and Nb35 were further refined by manual model building in COOT(Emsley et al. 2010) and real space refinement, as implemented in the Phenix software(Adams et al. 2010). The peptide C terminus, ECD and ECLs were modelled manually without ambiguity with the aid of ECD-focused map. Density of the linker (K130^ECD^ – S136^ECD^) and ICL3 (L399^ICL3^ – T343^ICL3^) was not clear and these residues were omitted from the final model. The AHD of taspoglutide complex was modelled according to the AHD-focused map with side chain of 31 residues deleted. However, the AHD of the semaglutide complex was not able to be modelled due to the low resolution of this region. The final models were subjected to global refinement and comprehensive validation. The cryo-EM data collection, refinement and validation statistics are reported in Table S1.

### Model residue interaction analysis

Interactions in the PDB files between the bound peptide and the receptor were analyzed using the “Dimplot” module within the Ligplot+ program (v2.2)(Laskowski and Swindells 2011). Hydrogen bonds were additionally analysed using the UCSF Chimera package(Pettersen et al. 2004), with relaxed distance and angle criteria (0.4 Å and 20 degree tolerance, respectively).

### CryoEM dynamics analysis

3D variability analysis implemented in cryoSPARC (v2.9)(Punjani et al. 2017) was performed to understand and visualize the dynamics in GLP-1R complexes, as previously described for analysis of the dynamics of adrenomedullin receptors(Liang, Belousoff, Fletcher, et al. 2020). The particle stack of semaglutide and taspoglutide-GLP-1R-Gs complexes from RELION consensus refinement were imported into the cryoSPARC environment. 3D refinement was performed, using a low pass filtered RELION consensus map as an initial model and a generous mask created in cryoSPARC as default. 3D variability of these GLP-1R complexes was analysed in 5 motions and the 20 volume frame data in each motion was generated in cryoSPARC(Punjani et al. 2017). Output files were visualized in UCSF ChimeraX volume series and captured as movies(Goddard et al. 2018).

The backbone of the both peptide-bound GLP-1R was rigid body fitted into the two maps at the extremes of the mode (frame000 and frame019) of component 1 and further refined by manual model building in COOT. The ECD of taspoglutide-bound receptor was modelled by rigid body fit and manual model building into the two extreme maps (frame000 and frame019) of component 4. The consensus ECD of semaglutide-bound receptor was rigid body fitted into frame000 map of component 4.

### Stable cell lines generation

The wild-type (WT) and mutant GLP-1R constructs were integrated into FlpIn-CHO cells using FlpIn Gateway technology system (Invitrogen). Stable CHOFlpIn expression cell lines were selected using 600 μg/ml hygromyocin B, and maintained in DMEM supplemented with 5% (V/V) FBS (Invitrogen) at 37°C in 5% CO_2_.

### cAMP accumulation assay

cAMP accumulation assay was performed as described previously(Hager et al. 2017). CHOFlpIn WT or mutant GLP-1R cells were seeded at a density of 30,000 cells per well into a 96-well plate and incubated overnight at 37°C in 5% CO_2_. All values were converted to cAMP concentration using cAMP standard curve performed parallel and data were subsequently normalized to the response of 100μM forskolin in each cell line, and then normalized to the WT for each peptide agonist.

### Data analysis

Pharmacological data were analysed using a 3-parameter logistic fit in Prism (v8.2, GraphPad). Data were analysed using a 3-parameter logistic fit in Prism and assessed for differences in fitted parameters from the parental construct at 95% confidence intervals. Differences in globally fitted curves were also assessed using an extra sum of squares F test at P<0.05, with post-hoc assessment of individual fitted parameters where curves were statistically different.

### Data availability

The atomic coordinates and the cryo-EM density maps generated during this study are available at the protein databank (https://www.rcsb.org) and the electron microscopy databank (https://www.ebi.ac.uk/pdbe/emdb) under accession numbers XXXX and XXXX, and EMDB entry ID EMD-XXXXX and ID EMD-XXXXX for the semaglutide and taspogltuide complexes, respectively.

## SUPPLEMENTAL INFORMATION

**Video S1. 3D variability analysis of semaglutide- and taspoglutide-bound GLP-1R-G**_**s**_ **complexes**. CryoSPARC variability analysis performed on the GLP-1R-G_s_ complexes in the presence of different bound agonists. Transition 1: Principal component 1; Transition 2: Principal component 2; Transition 3; Principal component 3.

**Video S2. 3D variability analysis reveals differences in the conformational dynamics of the GLP-1R ECD and Gs AHD in the presence of semaglutide versus taspoglutide**. Transition 1: CryoSPARC 3D variability analysis principal component 4. This illustrates differences in the dynamics of the receptor ECD. Transition 2: CryoSPARC 3D variability analysis of principal component 5 for the semaglutide-bound complex and principal component 1 for the taspoglutide-bound complex. These illustrate the differences in the dynamics of the Gα AHD. Transition 3: morphs between the three major classes of location of the Gs AHD for the semaglutide and taspoglutide bound GLP-1R-Gs complexes determined from focused 3D classification in RELION.

## REFERENCES

Adams, P. D., P. V. Afonine, G. Bunkoczi, V. B. Chen, I. W. Davis, N. Echols, J. J. Headd, L. W. Hung, G. J. Kapral, R. W. Grosse-Kunstleve, A. J. McCoy, N. W. Moriarty, R. Oeffner, R. J. Read, D. C. Richardson, J. S. Richardson, T. C. Terwilliger, and P. H. Zwart. 2010. “PHENIX: a comprehensive Python-based system for macromolecular structure solution.” Acta Crystallogr D Biol Crystallogr 66 (Pt 2):213–21. doi: 10.1107/S0907444909052925.

Adelhorst, K., B. B. Hedegaard, L. B. Knudsen, and O. Kirk. 1994. “Structure-activity studies of glucagon-like peptide-1.” J Biol Chem 269 (9):6275–8.

Ahmann, A. J., M. Capehorn, G. Charpentier, F. Dotta, E. Henkel, I. Lingvay, A. G. Holst, M. P. Annett, and V. R. Aroda. 2018. “Efficacy and Safety of Once-Weekly Semaglutide Versus Exenatide ER in Subjects With Type 2 Diabetes (SUSTAIN 3): A 56-Week, Open-Label, Randomized Clinical Trial.” Diabetes Care 41 (2):258–266. doi: 10.2337/dc17-0417.

Chan, K. Y., L. G. Trabuco, E. Schreiner, and K. Schulten. 2012. “Cryo-electron microscopy modeling by the molecular dynamics flexible fitting method.” Biopolymers 97 (9):678–86. doi: 10.1002/bip.22042.

Day, J. W., P. Li, J. T. Patterson, J. Chabenne, M. D. Chabenne, V. M. Gelfanov, and R. D. Dimarchi. 2011. “Charge inversion at position 68 of the glucagon and glucagon-like peptide-1 receptors supports selectivity in hormone action.” J Pept Sci 17 (3):218–25. doi: 10.1002/psc.1317.

Deacon, C. F., L. B. Knudsen, K. Madsen, F. C. Wiberg, O. Jacobsen, and J. J. Holst. 1998. “Dipeptidyl peptidase IV resistant analogues of glucagon-like peptide-1 which have extended metabolic stability and improved biological activity.” Diabetologia 41 (3):271–8. doi: 10.1007/s001250050903.

Dods, R. L., and D. Donnelly. 2015. “The peptide agonist-binding site of the glucagon-like peptide-1 (GLP-1) receptor based on site-directed mutagenesis and knowledge-based modelling.” Biosci Rep 36 (1):e00285. doi: 10.1042/BSR20150253.

Dong, M., G. Deganutti, S. J. Piper, Y. L. Liang, M. Khoshouei, M. J. Belousoff, K. G. Harikumar, C. A. Reynolds, A. Glukhova, S. G. B. Furness, A. Christopoulos, R. Danev, D. Wootten, P. M. Sexton, and L. J. Miller. 2020. “Structure and dynamics of the active Gs-coupled human secretin receptor.” Nat Commun 11 (1):4137. doi: 10.1038/s41467-020-17791-4.

Draper-Joyce, C. J., M. Khoshouei, D. M. Thal, Y. L. Liang, A. T. N. Nguyen, S. G. B. Furness, H. Venugopal, J. A. Baltos, J. M. Plitzko, R. Danev, W. Baumeister, L. T. May, D. Wootten, P. M. Sexton, A. Glukhova, and A. Christopoulos. 2018. “Structure of the adenosine-bound human adenosine A1 receptor-Gi complex.” Nature 558 (7711):559–563. doi: 10.1038/s41586-018-0236-6.

Emsley, P., B. Lohkamp, W. G. Scott, and K. Cowtan. 2010. “Features and development of Coot.” Acta Crystallogr D Biol Crystallogr 66 (Pt 4):486–501. doi: 10.1107/S0907444910007493.

Fletcher, M. M., M. L. Halls, P. Zhao, L. Clydesdale, A. Christopoulos, P. M. Sexton, and D. Wootten. 2018. “Glucagon-like peptide-1 receptor internalisation controls spatiotemporal signalling mediated by biased agonists.” Biochem Pharmacol 156:406–419. doi: 10.1016/j.bcp.2018.09.003.

Fletcher, M. M. 2019 The spatiotemporal control of signalling and trafficking of the GLP-1R. Monash University thesis. https://doi.org/10.26180/5c4fd15a05152

Furness, S. G. B., Y. L. Liang, C. J. Nowell, M. L. Halls, P. J. Wookey, E. Dal Maso, A. Inoue, A. Christopoulos, D. Wootten, and P. M. Sexton. 2016. “Ligand-Dependent Modulation of G Protein Conformation Alters Drug Efficacy.” Cell 167 (3):739–749 e11. doi: 10.1016/j.cell.2016.09.021.

Garcia-Nafria, J., and C. G. Tate. 2019. “Cryo-EM structures of GPCRs coupled to Gs, Gi and Go.” Mol Cell Endocrinol 488:1–13. doi: 10.1016/j.mce.2019.02.006.

Goddard, T. D., C. C. Huang, E. C. Meng, E. F. Pettersen, G. S. Couch, J. H. Morris, and T. E. Ferrin. 2018. “UCSF ChimeraX: Meeting modern challenges in visualization and analysis.” Protein Sci 27 (1):14–25. doi: 10.1002/pro.3235.

Graaf, Cd, D. Donnelly, D. Wootten, J. Lau, P. M. Sexton, L. J. Miller, J. M. Ahn, J. Liao, M. M. Fletcher, D. Yang, A. J. Brown, C. Zhou, J. Deng, and M. W. Wang. 2016. “Glucagon-Like Peptide-1 and Its Class B G Protein-Coupled Receptors: A Long March to Therapeutic Successes.” Pharmacol Rev 68 (4):954–1013. doi: 10.1124/pr.115.011395.

Hager, M. V., L. Clydesdale, S. H. Gellman, P. M. Sexton, and D. Wootten. 2017. “Characterization of signal bias at the GLP-1 receptor induced by backbone modification of GLP-1.” Biochem Pharmacol 136:99–108. doi: 10.1016/j.bcp.2017.03.018.

Hauser, A. S., M. M. Attwood, M. Rask-Andersen, H. B. Schioth, and D. E. Gloriam. 2017. “Trends in GPCR drug discovery: new agents, targets and indications.” Nat Rev Drug Discov 16 (12):829–842. doi: 10.1038/nrd.2017.178.

Henry, R. R., S. Mudaliar, L. Kanitra, M. Woloschak, R. Balena, and T. Emerge 3 Study Group. 2012. “Efficacy and safety of taspoglutide in patients with type 2 diabetes inadequately controlled with metformin plus pioglitazone over 24 weeks: T-Emerge 3 trial.” J Clin Endocrinol Metab 97 (7):2370–9. doi: 10.1210/jc.2011-3253.

Jones, B., T. Buenaventura, N. Kanda, P. Chabosseau, B. M. Owen, R. Scott, R. Goldin, N. Angkathunyakul, I. R. Correa, Jr., D. Bosco, P. R. Johnson, L. Piemonti, P. Marchetti, A. M. J. Shapiro, B. J. Cochran, A. C. Hanyaloglu, A. Inoue, T. Tan, G. A. Rutter, A. Tomas, and S. R. Bloom. 2018. “Targeting GLP-1 receptor trafficking to improve agonist efficacy.” Nat Commun 9 (1):1602. doi: 10.1038/s41467-018-03941-2.

Koole, C., D. Wootten, J. Simms, L. J. Miller, A. Christopoulos, and P. M. Sexton. 2012. “Second extracellular loop of human glucagon-like peptide-1 receptor (GLP-1R) has a critical role in GLP-1 peptide binding and receptor activation.” J Biol Chem 287 (6):3642–58. doi: 10.1074/jbc.M111.309328.

Koole, C., D. Wootten, J. Simms, C. Valant, R. Sridhar, O. L. Woodman, L. J. Miller, R. J. Summers, A. Christopoulos, and P. M. Sexton. 2010. “Allosteric ligands of the glucagon-like peptide 1 receptor (GLP-1R) differentially modulate endogenous and exogenous peptide responses in a pathway-selective manner: implications for drug screening.” Mol Pharmacol 78 (3):456–65. doi: 10.1124/mol.110.065664.

Laskowski, R. A., and M. B. Swindells. 2011. “LigPlot+: multiple ligand-protein interaction diagrams for drug discovery.” J Chem Inf Model 51 (10):2778–86. doi: 10.1021/ci200227u.

Lau, C., M. J. Hunter, A. Stewart, E. Perozo, and J. I. Vandenberg. 2018. “Never at rest: insights into the conformational dynamics of ion channels from cryo-electron microscopy.” J Physiol 596 (7):1107–1119. doi: 10.1113/JP274888.

Lau, J., P. Bloch, L. Schaffer, I. Pettersson, J. Spetzler, J. Kofoed, K. Madsen, L. B. Knudsen, J. McGuire, D. B. Steensgaard, H. M. Strauss, D. X. Gram, S. M. Knudsen, F. S. Nielsen, P. Thygesen, S. Reedtz-Runge, and T. Kruse. 2015. “Discovery of the Once-Weekly Glucagon-Like Peptide-1 (GLP-1) Analogue Semaglutide.” J Med Chem 58 (18):7370–80. doi: 10.1021/acs.jmedchem.5b00726.

Lei, S., L. Clydesdale, A. Dai, X. Cai, Y. Feng, D. Yang, Y. L. Liang, C. Koole, P. Zhao, T. Coudrat, A. Christopoulos, M. W. Wang, D. Wootten, and P. M. Sexton. 2018. “Two distinct domains of the glucagon-like peptide-1 receptor control peptide-mediated biased agonism.” J Biol Chem 293 (24):9370–9387. doi: 10.1074/jbc.RA118.003278.

Liang, Y. L., M. J. Belousoff, M. M. Fletcher, X. Zhang, M. Khoshouei, G. Deganutti, C. Koole, S. G. B. Furness, L. J. Miller, D. L. Hay, A. Christopoulos, C. A. Reynolds, R. Danev, D. Wootten, and P. M. Sexton. 2020. “Structure and Dynamics of Adrenomedullin Receptors AM1 and AM2 Reveal Key Mechanisms in the Control of Receptor Phenotype by Receptor Activity-Modifying Proteins.” ACS Pharmacol Transl Sci 3 (2):263–284. doi: 10.1021/acsptsci.9b00080.

Liang, Y. L., M. J. Belousoff, P. Zhao, C. Koole, M. M. Fletcher, T. T. Truong, V. Julita, G. Christopoulos, H. E. Xu, Y. Zhang, M. Khoshouei, A. Christopoulos, R. Danev, P. M. Sexton, and D. Wootten. 2020. “Toward a Structural Understanding of Class B GPCR Peptide Binding and Activation.” Mol Cell 77 (3):656–668 e5. doi: 10.1016/j.molcel.2020.01.012.

Liang, Y. L., M. Khoshouei, G. Deganutti, A. Glukhova, C. Koole, T. S. Peat, M. Radjainia, J. M. Plitzko, W. Baumeister, L. J. Miller, D. L. Hay, A. Christopoulos, C. A. Reynolds, D. Wootten, and P. M. Sexton. 2018. “Cryo-EM structure of the active, Gs-protein complexed, human CGRP receptor.” Nature 561 (7724):492–497. doi: 10.1038/s41586-018-0535-y.

Liang, Y. L., M. Khoshouei, A. Glukhova, S. G. B. Furness, P. Zhao, L. Clydesdale, C. Koole, T. T. Truong, D. M. Thal, S. Lei, M. Radjainia, R. Danev, W. Baumeister, M. W. Wang, L. J. Miller, A. Christopoulos, P. M. Sexton, and D. Wootten. 2018. “Phase-plate cryo-EM structure of a biased agonist-bound human GLP-1 receptor-Gs complex.” Nature 555 (7694):121–125. doi: 10.1038/nature25773.

Liang, Y. L., M. Khoshouei, M. Radjainia, Y. Zhang, A. Glukhova, J. Tarrasch, D. M. Thal, S. G. B. Furness, G. Christopoulos, T. Coudrat, R. Danev, W. Baumeister, L. J. Miller, A. Christopoulos, B. K. Kobilka, D. Wootten, G. Skiniotis, and P. M. Sexton. 2017. “Phase-plate cryo-EM structure of a class B GPCR-G-protein complex.” Nature 546 (7656):118–123. doi: 10.1038/nature22327.

Liang, Yi-Lynn, Peishen Zhao, Christopher Draper-Joyce, Jo-Anne Baltos, Alisa Glukhova, Tin T. Truong, Lauren T. May, Arthur Christopoulos, Denise Wootten, Patrick M. Sexton, and Sebastian G. B. Furness. 2018. “Dominant Negative G Proteins Enhance Formation and Purification of Agonist-GPCR-G Protein Complexes for Structure Determination.” ACS Pharmacology & Translational Science 1 (1):12–20. doi: 10.1021/acsptsci.8b00017.

Maeda, S., Q. Qu, M. J. Robertson, G. Skiniotis, and B. K. Kobilka. 2019. “Structures of the M1 and M2 muscarinic acetylcholine receptor/G-protein complexes.” Science 364 (6440):552–557. doi: 10.1126/science.aaw5188.

Marso, S. P., S. C. Bain, A. Consoli, F. G. Eliaschewitz, E. Jodar, L. A. Leiter, I. Lingvay, J. Rosenstock, J. Seufert, M. L. Warren, V. Woo, O. Hansen, A. G. Holst, J. Pettersson, T. Vilsboll, and Sustain-Investigators. 2016. “Semaglutide and Cardiovascular Outcomes in Patients with Type 2 Diabetes.” N Engl J Med 375 (19):1834–1844. doi: 10.1056/NEJMoa1607141.

Murata, K., and M. Wolf. 2018. “Cryo-electron microscopy for structural analysis of dynamic biological macromolecules.” Biochim Biophys Acta Gen Subj 1862 (2):324–334. doi: 10.1016/j.bbagen.2017.07.020.

O’Neil, P. M., A. L. Birkenfeld, B. McGowan, O. Mosenzon, S. D. Pedersen, S. Wharton, C. G. Carson, C. H. Jepsen, M. Kabisch, and J. P. H. Wilding. 2018. “Efficacy and safety of semaglutide compared with liraglutide and placebo for weight loss in patients with obesity: a randomised, double-blind, placebo and active controlled, dose-ranging, phase 2 trial.” Lancet 392 (10148):637–649. doi: 10.1016/S0140-6736(18)31773-2.

Pettersen, E. F., T. D. Goddard, C. C. Huang, G. S. Couch, D. M. Greenblatt, E. C. Meng, and T. E. Ferrin. 2004. “UCSF Chimera--a visualization system for exploratory research and analysis.” J Comput Chem 25 (13):1605–12. doi: 10.1002/jcc.20084.

Pickford, P., M. Lucey, Z. Fang, S. Bitsi, J. B. de la Serna, J. Broichhagen, D. J. Hodson, J. Minnion, G. A. Rutter, S. R. Bloom, A. Tomas, and B. Jones. 2020. “Signalling, trafficking and glucoregulatory properties of glucagon-like peptide-1 receptor agonists exendin-4 and lixisenatide.” Br J Pharmacol. doi: 10.1111/bph.15134.

Pratley, R. E., V. R. Aroda, I. Lingvay, J. Ludemann, C. Andreassen, A. Navarria, A. Viljoen, and Sustain investigators. 2018. “Semaglutide versus dulaglutide once weekly in patients with type 2 diabetes (SUSTAIN 7): a randomised, open-label, phase 3b trial.” Lancet Diabetes Endocrinol 6 (4):275–286. doi: 10.1016/S2213-8587(18)30024-X.

Punjani, A., J. L. Rubinstein, D. J. Fleet, and M. A. Brubaker. 2017. “cryoSPARC: algorithms for rapid unsupervised cryo-EM structure determination.” Nat Methods 14 (3):290–296. doi: 10.1038/nmeth.4169.

Punjani, Ali, and David J. Fleet. 2020. “3D Variability Analysis: Directly resolving continuous flexibility and discrete heterogeneity from single particle cryo-EM images.” bioRxiv:2020.04.08.032466. doi: 10.1101/2020.04.08.032466.

Qiao, A., S. Han, X. Li, Z. Li, P. Zhao, A. Dai, R. Chang, L. Tai, Q. Tan, X. Chu, L. Ma, T. S. Thorsen, S. Reedtz-Runge, D. Yang, M. W. Wang, P. M. Sexton, D. Wootten, F. Sun, Q. Zhao, and B. Wu. 2020. “Structural basis of Gs and Gi recognition by the human glucagon receptor.” Science 367 (6484):1346–1352. doi: 10.1126/science.aaz5346.

Rasmussen, S. G., B. T. DeVree, Y. Zou, A. C. Kruse, K. Y. Chung, T. S. Kobilka, F. S. Thian, P. S. Chae, E. Pardon, D. Calinski, J. M. Mathiesen, S. T. Shah, J. A. Lyons, M. Caffrey, S. H. Gellman, J. Steyaert, G. Skiniotis, W. I. Weis, R. K. Sunahara, and B. K. Kobilka. 2011. “Crystal structure of the beta2 adrenergic receptor-Gs protein complex.” Nature 477 (7366):549–55. doi: 10.1038/nature10361.

Rosenstock, J., B. Balas, B. Charbonnel, G. B. Bolli, M. Boldrin, R. Ratner, R. Balena, and T. emerge 2 Study Group. 2013. “The fate of taspoglutide, a weekly GLP-1 receptor agonist, versus twice-daily exenatide for type 2 diabetes: the T-emerge 2 trial.” Diabetes Care 36 (3):498–504. doi: 10.2337/dc12-0709.

Schorb, M., I. Haberbosch, W. J. H. Hagen, Y. Schwab, and D. N. Mastronarde. 2019. “Software tools for automated transmission electron microscopy.” Nat Methods 16 (6):471–477. doi: 10.1038/s41592-019-0396-9.

Sebokova, E., A. D. Christ, H. Wang, S. Sewing, J. Z. Dong, J. Taylor, M. A. Cawthorne, and M. D. Culler. 2010. “Taspoglutide, an analog of human glucagon-like Peptide-1 with enhanced stability and in vivo potency.” Endocrinology 151 (6):2474–82. doi: 10.1210/en.2009-1459.

Sun, Feng, Sanbao Chai, Kai Yu, Xiaochi Quan, Zhirong Yang, Shanshan Wu, Yuan Zhang, Linong Ji, Jun Wang, and Luwen Shi. 2014. “Gastrointestinal Adverse Events of Glucagon-Like Peptide- 1 Receptor Agonists in Patients with Type 2 Diabetes: A Systematic Review and Network Meta-Analysis.” Diabetes Technology & Therapeutics 17 (1):35–42. doi: 10.1089/dia.2014.0188.

Tsoukas, George, Stephen Bain, Eiichi Araki, Cyrus Desouza, Satish Garg, Ludger Rose, Eirik Bergan, Julie Karsbøl, and Hans Devries. 2017. “124 - Semaglutide Reduces HbA1c Across Baseline HbA1c Subgroups Across SUSTAIN 1–5 Clinical Trials.” Canadian Journal of Diabetes 41 (5, Supplement):S50. doi: https://doi.org/10.1016/j.jcjd.2017.08.132.

Underwood, C. R., P. Garibay, L. B. Knudsen, S. Hastrup, G. H. Peters, R. Rudolph, and S. Reedtz-Runge. 2010. “Crystal structure of glucagon-like peptide-1 in complex with the extracellular domain of the glucagon-like peptide-1 receptor.” J Biol Chem 285 (1):723–30. doi: 10.1074/jbc.M109.033829.

Wagner, T., F. Merino, M. Stabrin, T. Moriya, C. Antoni, A. Apelbaum, P. Hagel, O. Sitsel, T. Raisch, D. Prumbaum, D. Quentin, D. Roderer, S. Tacke, B. Siebolds, E. Schubert, T. R. Shaikh, P. Lill, C. Gatsogiannis, and S. Raunser. 2019. “SPHIRE-crYOLO is a fast and accurate fully automated particle picker for cryo-EM.” Commun Biol 2:218. doi: 10.1038/s42003-019-0437-z.

Westfield, G. H., S. G. Rasmussen, M. Su, S. Dutta, B. T. DeVree, K. Y. Chung, D. Calinski, G. Velez-Ruiz, A. N. Oleskie, E. Pardon, P. S. Chae, T. Liu, S. Li, V. L. Woods, Jr., J. Steyaert, B. K. Kobilka, R. K. Sunahara, and G. Skiniotis. 2011. “Structural flexibility of the G alpha s alpha-helical domain in the beta2-adrenoceptor Gs complex.” Proc Natl Acad Sci U S A 108 (38):16086–91. doi: 10.1073/pnas.1113645108.

Wilmen, Andreas, Brigitte Van Eyll, Burkhard Göke, and Rüdiger Göke. 1997. “Five Out of Six Tryptophan Residues in the N-Terminal Extracellular Domain of the Rat GLP-1 Receptor Are Essential for its Ability to Bind GLP-1.” Peptides 18 (2):301–305. doi: https://doi.org/10.1016/S0196-9781(96)00321-X.

Wootten, D., C. A. Reynolds, C. Koole, K. J. Smith, J. C. Mobarec, J. Simms, T. Quon, T. Coudrat, S. G. Furness, L. J. Miller, A. Christopoulos, and P. M. Sexton. 2016. “A Hydrogen-Bonded Polar Network in the Core of the Glucagon-Like Peptide-1 Receptor Is a Fulcrum for Biased Agonism: Lessons from Class B Crystal Structures.” Mol Pharmacol 89 (3):335–47. doi: 10.1124/mol.115.101246.

Wootten, D., C. A. Reynolds, K. J. Smith, J. C. Mobarec, C. Koole, E. E. Savage, K. Pabreja, J. Simms, R. Sridhar, S. G. B. Furness, M. Liu, P. E. Thompson, L. J. Miller, A. Christopoulos, and P. M. Sexton. 2016. “The Extracellular Surface of the GLP-1 Receptor Is a Molecular Trigger for Biased Agonism.” Cell 165 (7):1632–1643. doi: 10.1016/j.cell.2016.05.023.

Wootten, D., J. Simms, L. J. Miller, A. Christopoulos, and P. M. Sexton. 2013. “Polar transmembrane interactions drive formation of ligand-specific and signal pathway-biased family B G protein-coupled receptor conformations.” Proceedings of the National Academy of Sciences 110 (13):5211–5216. doi: 10.1073/pnas.1221585110.

Yang, D., C. de Graaf, L. Yang, G. Song, A. Dai, X. Cai, Y. Feng, S. Reedtz-Runge, M. A. Hanson, H. Yang, H. Jiang, R. C. Stevens, and M. W. Wang. 2016. “Structural Determinants of Binding the Seven-transmembrane Domain of the Glucagon-like Peptide-1 Receptor (GLP-1R).” J Biol Chem 291 (25):12991–3004. doi: 10.1074/jbc.M116.721977.

Yin, Y., X. E. Zhou, L. Hou, L. H. Zhao, B. Liu, G. Wang, Y. Jiang, K. Melcher, and H. E. Xu. 2016. “An intrinsic agonist mechanism for activation of glucagon-like peptide-1 receptor by its extracellular domain.” Cell Discov 2:16042. doi: 10.1038/celldisc.2016.42.

Zhang, K. 2016. “Gctf: Real-time CTF determination and correction.” J Struct Biol 193 (1):1–12. doi: 10.1016/j.jsb.2015.11.003.

Zhang, X., M. J. Belousoff, P. Zhao, A. J. Kooistra, T. T. Truong, S. Y. Ang, C. R. Underwood, T. Egebjerg, P. Senel, G. D. Stewart, Y. L. Liang, A. Glukhova, H. Venugopal, A. Christopoulos, S. G. B. Furness, L. J. Miller, S. Reedtz-Runge, C. J. Langmead, D. E. Gloriam, R. Danev, P. M. Sexton, and D. Wootten. 2020. “Differential GLP-1R Binding and Activation by Peptide and Non-peptide Agonists.” Mol Cell. doi: 10.1016/j.molcel.2020.09.020.

Zhao, L. H., S. Ma, I. Sutkeviciute, D. D. Shen, X. E. Zhou, P. W. de Waal, C. Y. Li, Y. Kang, L. J. Clark, F. G. Jean-Alphonse, A. D. White, D. Yang, A. Dai, X. Cai, J. Chen, C. Li, Y. Jiang, T. Watanabe, T. J. Gardella, K. Melcher, M. W. Wang, J. P. Vilardaga, H. E. Xu, and Y. Zhang. 2019. “Structure and dynamics of the active human parathyroid hormone receptor-1.” Science 364 (6436):148–153. doi: 10.1126/science.aav7942.

Zhao, P., Y. L. Liang, M. J. Belousoff, G. Deganutti, M. M. Fletcher, F. S. Willard, M. G. Bell, M. E. Christe, K. W. Sloop, A. Inoue, T. T. Truong, L. Clydesdale, S. G. B. Furness, A. Christopoulos, M. W. Wang, L. J. Miller, C. A. Reynolds, R. Danev, P. M. Sexton, and D. Wootten. 2020. “Activation of the GLP-1 receptor by a non-peptidic agonist.” Nature 577 (7790):432–436. doi: 10.1038/s41586-019-1902-z.

Zheng, S. Q., E. Palovcak, J. P. Armache, K. A. Verba, Y. Cheng, and D. A. Agard. 2017. “MotionCor2: anisotropic correction of beam-induced motion for improved cryo-electron microscopy.” Nat Methods 14 (4):331–332. doi: 10.1038/nmeth.4193.

Zivanov, J., T. Nakane, B. O. Forsberg, D. Kimanius, W. J. Hagen, E. Lindahl, and S. H. Scheres. 2018. “New tools for automated high-resolution cryo-EM structure determination in RELION-3.” Elife 7. doi: 10.7554/eLife.42166.

