## Supplemental Tables and Figures for "Structure and dynamics of semaglutide and taspoglutide bound GLP-1R-Gs complexes"

**Table S1. Data collection and refinement statistics.** *Related to Figures 1, S1 and S2.*

| **Data Collection** | Sema:GLP-1R:DNGs:Nb35 | Taspo:GLP-1R:DNGs:Nb35 |
|---|---|---|
| Micrographs | 7488 | 8759 |
| Electron dose ($e^-/A^2$) | 66.5 | 66.5 |
| Voltage (kV) | 300 | 300 |
| Pixel size (Å) | 0.83 | 1.207 |
| Defocus range (µm) | 0.5-1.5 | 0.5-1.5 |
| Symmetry imposed | C1 | C1 |
| Particles (final map) | 886,738 | 625,241 |
| Resolution (0.143 FSC)(Å) | 2.5 | 2.5 |
| **Refinement** | | |
| $CC_{map\_model}$ | | |
| Map sharpening B factor (Å$^2$) | -55 | -70 |
| **Model Quality** | | |
| R.m.s. deviations | | |
| Bond length (Å) | 0.007 | 0.008 |
| Bond angles (°) | 0.850 | 0.935 |
| Ramachandran | | |
| Favoured (%) | 96.95 | 96.51 |
| Outliers (%) | 0 | 0 |
| Rotamer outliers | 0 | 0.28 |
| C-Beta deviations (%) | 0 | 0 |
| Clashscore | 7.52 | 8.06 |
| MolProbity score | 1.59 | 1.67 |

**Table S2: Interactions between the GLP-1R and semaglutide or taspoglutide.** *Related to Figures 2, 3 and S4.*

| Peptide | Semaglutide:GLP-1R:DNGs | Taspoglutide:GLP-1R:DNGs | Interactions |
|---|---|---|---|
| His7 | Q234$^{3.37}$<br>*Water | Q234$^{3.37}$<br>*Water | Hydrogen bond |
| | Q234$^{3.37}$<br>V237$^{3.40}$<br>W306$^{5.36}$<br>I309$^{5.39}$<br>R310$^{5.40}$<br>I313$^{5.43}$ | Q234$^{3.37}$<br>V237$^{3.40}$<br>W306$^{5.36}$<br>I309$^{5.39}$<br>R310$^{5.40}$<br>I313$^{5.43}$ | Hydrophobic interactions |
| Aib8 | Y241$^{3.44}$<br>*K383$^{7.38}$<br>*E387$^{7.42}$ | | Hydrogen bond via water(s) |
| | L384$^{7.39}$<br>E387$^{7.42}$ | L384$^{7.39}$<br>E387$^{7.42}$ | Hydrophobic interactions |
| Glu9 | Y152$^{1.47}$<br>*R190$^{2.60}$<br>*T391$^{7.46}$ (3.5Å) | Y152$^{1.47}$<br>R190$^{2.60}$ | Hydrogen bond |
| | Y152$^{1.47}$<br>R190$^{2.60}$ | Y152$^{1.47}$<br>R190$^{2.60}$<br>K197$^{2.67}$<br>V237$^{3.40}$<br>L388$^{7.43}$ | Hydrophobic interactions |
| Gly10 | | *N300$^{ECL2}$ (bb, 3.6 Å) | Hydrogen bond |
| | | W306$^{ECL2}$ | Hydrophobic interaction |
| Thr11 | *D372$^{ECL3}$ (3.4Å) | D372$^{ECL3}$ | Hydrogen bond |
| | D372$^{ECL3}$<br>R380$^{7.35}$<br>L384$^{7.39}$ | D372$^{ECL3}$<br>R380$^{7.35}$<br>L384$^{7.39}$ | Hydrophobic interactions |
| Phe12 | L141$^{1.36}$<br>L144$^{1.39}$<br>Y148$^{1.43}$ | L141$^{1.36}$<br>L144$^{1.39}$<br>Y148$^{1.43}$<br>L388$^{7.43}$ | Hydrophobic interactions<br>π-π interactions |
| Thr13 | K197$^{2.67}$ | K197$^{2.67}$ | Hydrogen bond |
| | K197$^{2.67}$<br>L201$^{2.71}$<br>F230$^{3.33}$ | K197$^{2.67}$<br>L201$^{2.71}$<br>F230$^{3.33}$<br>M233$^{3.36}$ | Hydrophobic interactions |
| Ser14 | N300$^{ECL2}$ (bb) | N300$^{ECL2}$ (bb) | Hydrogen bond |
| | T298$^{ECL2}$<br>R299$^{ECL2}$<br>N300$^{ECL2}$ | T298$^{ECL2}$<br>R299$^{ECL2}$<br>N300$^{ECL2}$ | Hydrophobic interactions |
| Asp15 | R380$^{7.35}$ | R380$^{7.35}$ | Hydrogen bond/ionic |
| | R380$^{7.35}$ | R380$^{7.35}$<br>L384$^{7.39}$ | Hydrophobic interactions |

| | | | |
|---|---|---|---|
| Val16 | L141$^{1.36}$<br>L201$^{2.71}$ | L141$^{1.36}$ | Hydrophobic interactions |
| Ser17 | Y205$^{2.75}$<br>T298$^{ECL2}$<br>R299$^{ECL2}$ | Y205$^{2.75}$<br>T298$^{ECL2}$<br>R299$^{ECL2}$ | Hydrogen bond, |
| | T298$^{ECL2}$<br>R299$^{ECL2}$ | T298$^{ECL2}$<br>R299$^{ECL2}$ | Hydrophobic interactions |
| Ser18 | R299$^{ECL2}$ | R299$^{ECL2}$ | hydrophobic interactions |
| Tyr19 | E138$^{1.33}$ | E138$^{1.33}$ | Hydrogen bond |
| | E138$^{1.33}$ | E138$^{1.33}$ | Hydrophobic interactions |
| Leu20 | Y205$^{2.75}$ | Y205$^{2.75}$ | Hydrophobic interactions |
| Glu21 | S31$^{ECD}$<br>*R299$^{ECL2}$ | S31$^{ECD}$<br>L32$^{ECD}$ (bb)<br>Y205$^{2.75}$<br>R299$^{ECL2}$ | Hydrogen bond/Ionic interaction |
| | V30$^{ECD}$<br>S31$^{ECD}$<br>R299$^{ECL2}$ | V30$^{ECD}$<br>S31$^{ECD}$<br>L32$^{ECD}$<br>Y205$^{2.75}$<br>R299$^{ECL2}$ | Hydrophobic interactions |
| Ala24 | L32$^{ECD}$<br>Q210$^{ECL1}$ | L32$^{ECD}$<br>A209$^{ECL1}$ | Hydrophobic interactions |
| Ala25 | P90$^{ECD}$ | T35$^{ECD}$ | Hydrophobic interactions |
| Lys26 | | *E128$^{ECD}$ | Ionic interaction |
| Glu27 | W214$^{ECL1}$ | W214$^{ECL1}$ | Hydrophobic interactions |
| Phe28 | T35$^{ECD}$<br>W214$^{ECL1}$ | W39$^{ECD}$ | Hydrophobic interactions<br>π-π interactions |
| Ile29 | Y88$^{ECD}$<br>L89$^{ECD}$<br>P90$^{ECD}$<br>W91$^{ECD}$ | Y88$^{ECD}$<br>L89$^{ECD}$<br>P90$^{ECD}$<br>W91$^{ECD}$ | Hydrophobic interactions |
| Typ31 | W214$^{ECL1}$ | W214$^{ECL1}$ | π-π interactions |
| Leu32 | W39$^{ECD}$<br>E68$^{ECD}$ | W39$^{ECD}$<br>E68$^{ECD}$<br>Y88$^{ECD}$ | Hydrophobic interactions |
| Val33 | R121$^{ECD}$ | *R121$^{ECD}$ (3.4Å) | Hydrogen bond |
| | Y69$^{ECD}$<br>R121$^{ECD}$<br>L123$^{ECD}$ | Y69$^{ECD}$<br>R121$^{ECD}$ | Hydrophobic interactions |
| Gly$^{sema}$/Aib$^{taspo}$ 35 | | E68$^{ECD}$ | Hydrophobic interactions |
| R36$^{taspo}$ | W39$^{ECD}$<br>E68$^{ECD}$ | D67$^{ECD}$<br>E68$^{ECD}$ | Hydrophobic interactions |

Interactions in the PDB files between the peptide and the receptor were determined using Ligplot+. Hydrogen bonds were additionally determined using UCSF Chimera. Residues labelled with * are interactions not shown in Ligplot$^{+}$. bb, backbone.

**Video S1. 3D variability analysis of semaglutide- and taspoglutide- bound GLP-1R-$G_s$ complexes**. CryoSPARC variability analysis performed on the GLP-1R-$G_s$ complexes in the presence of different bound agonists. Transition 1: Principal component 1; Transition 2: Principal component 2; Transition 3; Principal component 3. *Related to Figures 5, 6 and S7.*

**Video S2. 3D variability analysis reveals differences in the conformational dynamics of the GLP-1R ECD and Gs AHD in the presence of semaglutide versus taspoglutide**. Transition 1: CryoSPARC 3D variability analysis principal component 4. This illustrates differences in the dynamics of the receptor ECD. Transition 2: CryoSPARC 3D variability analysis of principal component 5 for the semaglutide-bound complex and principal component 1 for the taspoglutide-bound complex. These illustrate the differences in the dynamics of the Gα AHD. Transition 3: morphs between the three major classes of location of the Gs AHD for the semaglutide and taspoglutide bound GLP-1R-Gs complexes determined from focused 3D classification in RELION. *Related to Figures 4, 5, 6 and S7.*

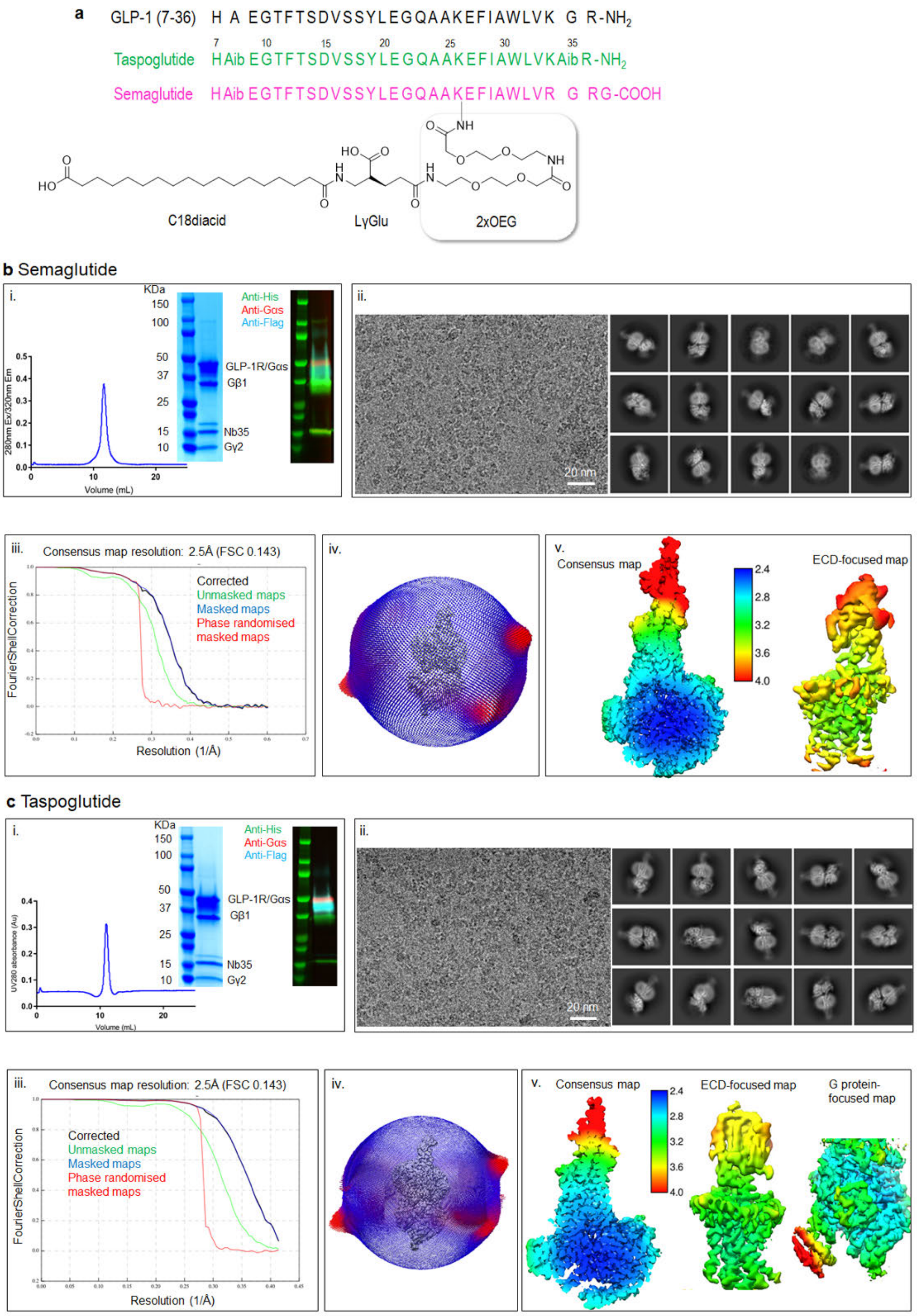


**Figure S1. Purification, cryo-EM imaging and processing of GLP-1R:G$_s$ complexes. a,** Amino acid sequence of GLP-1(7-36)NH$_2$, semaglutide and taspoglutide. Purification, cryo-EM imaging and processing of the semaglutide-bound complex (**b**) or taspoglutide-bound complex (**c**). Within each panel: (i) Size exclusion chromatography profile (left), SDS–PAGE/Coomassie blue stain and western blot of the complex showing all components (right). Anti-His antibody detects Gβ –His and Nb35–His (green), anti-Gs antibody detects Gαs (red) and anti-flag antibody detects Flag–GLP-1R–His. (ii) Exemplar micrograph (left) and 2D class averages of cryo-EM projections of the complex (right). (iii) Gold standard Fourier shell correlation 0.143 (FSC) curves for the final consensus maps and map validation from half maps, showing the overall nominal resolution. (iv) 3-D histogram representation of the Euler angle distribution of all the particles used in the reconstruction overlaid on the density map illustrated on the same coordinate axis. (v) Local resolution-filtered EM maps (consensus and ECD/G protein focused refinements) displaying local resolution (Å) coloured from highest resolution (dark blue) to lowest resolution (red). *Related to Figures 1, 2 and S2 and Table S1.*

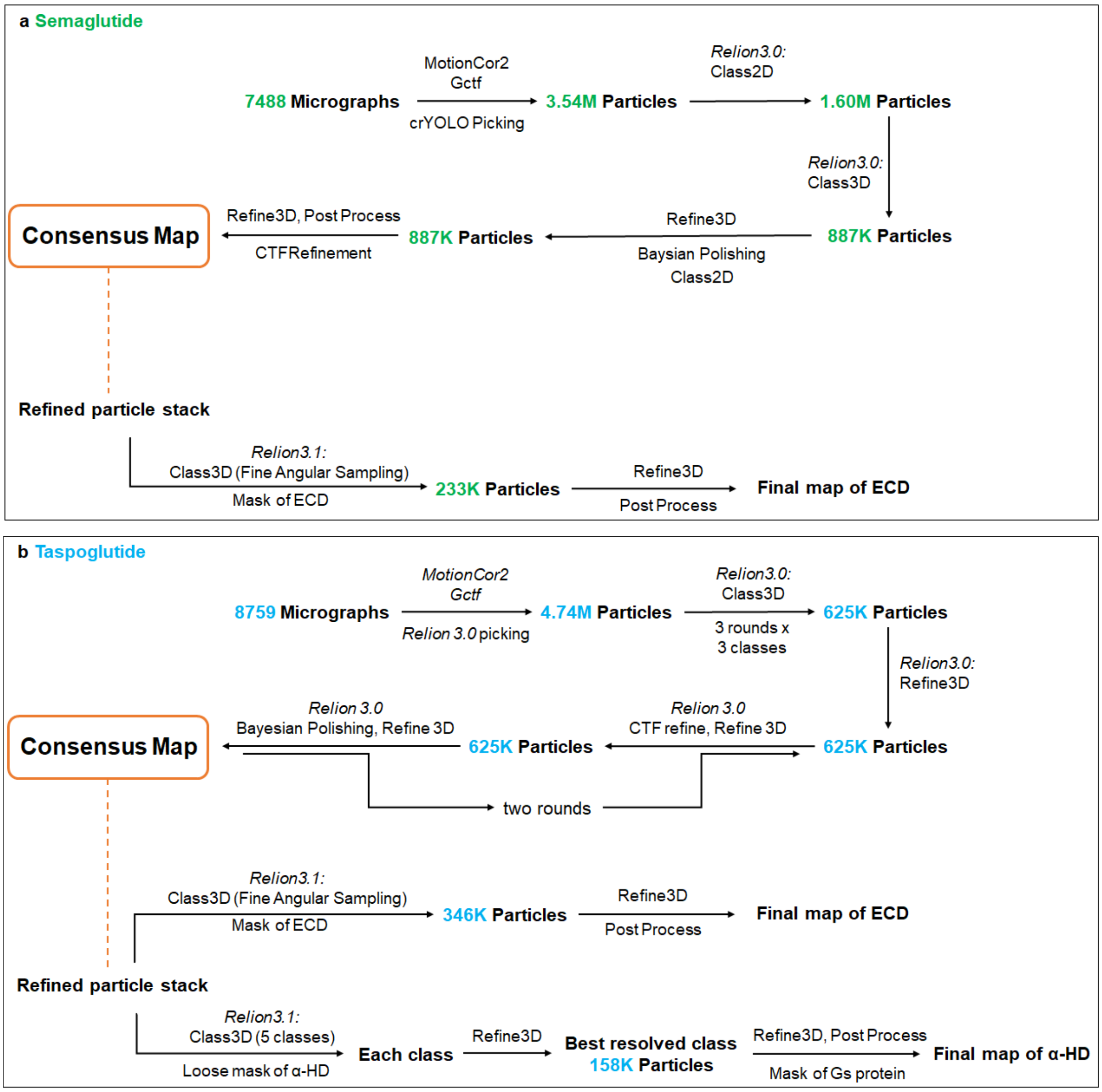


**Figure S2. Cryo-EM data processing workflow.** The number of micrographs or particles of semaglutide- (**a**) or taspoglutide- (**b**) bound complexes in each processing step are labelled in green or blue, respectively. The particles imported for local-focused refinement were from the particle stack for the consensus map construction as highlighted using rounded rectangle in orange. *Related to Figures 1 and S1 and Table S1.*

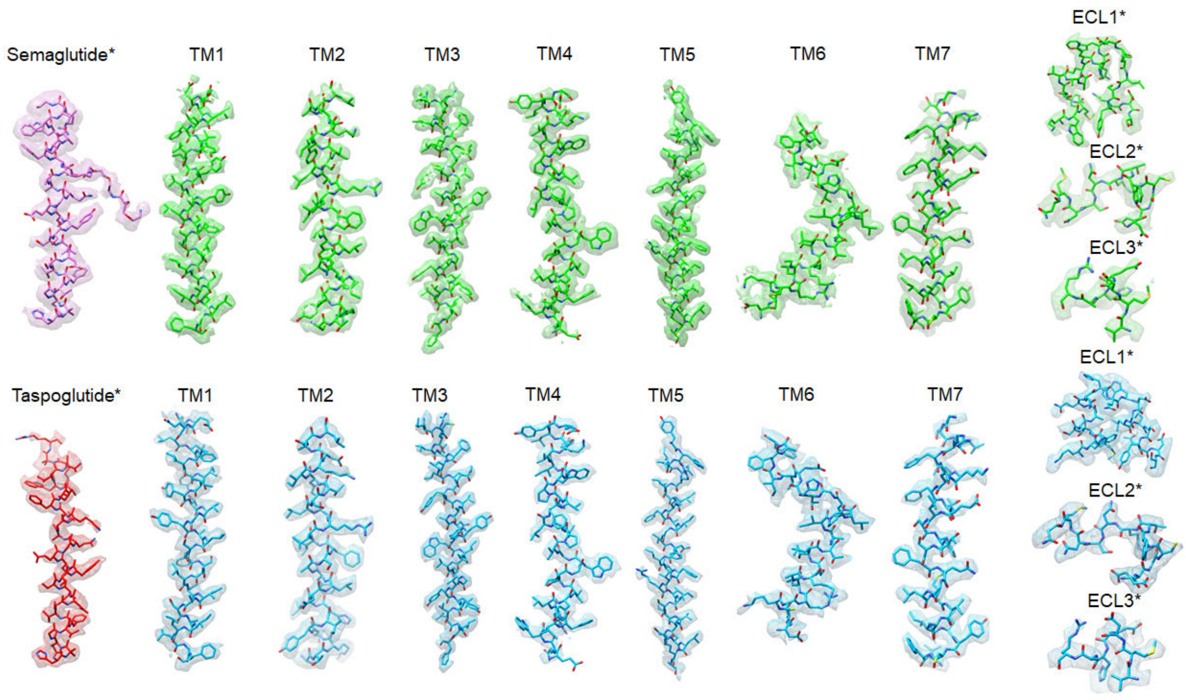


**Figure S3. Atomic models of the peptide, receptor TM helices and ECLs in the cryo-EM map.** The EM density map and model are shown for semaglutide (pink) and taspoglutide (red) and all seven TM helices, and ECLs of the GLP-1R when bound to each agonist (semaglutide bound GLP-1R: green; taspoglutide bound GLP-1R: blue). All residues are displayed as sticks coloured by heteroatom. The consensus map was used, with the exception of those labelled * where the ECD refined map is shown. *Related to Figures 1 and 2.*

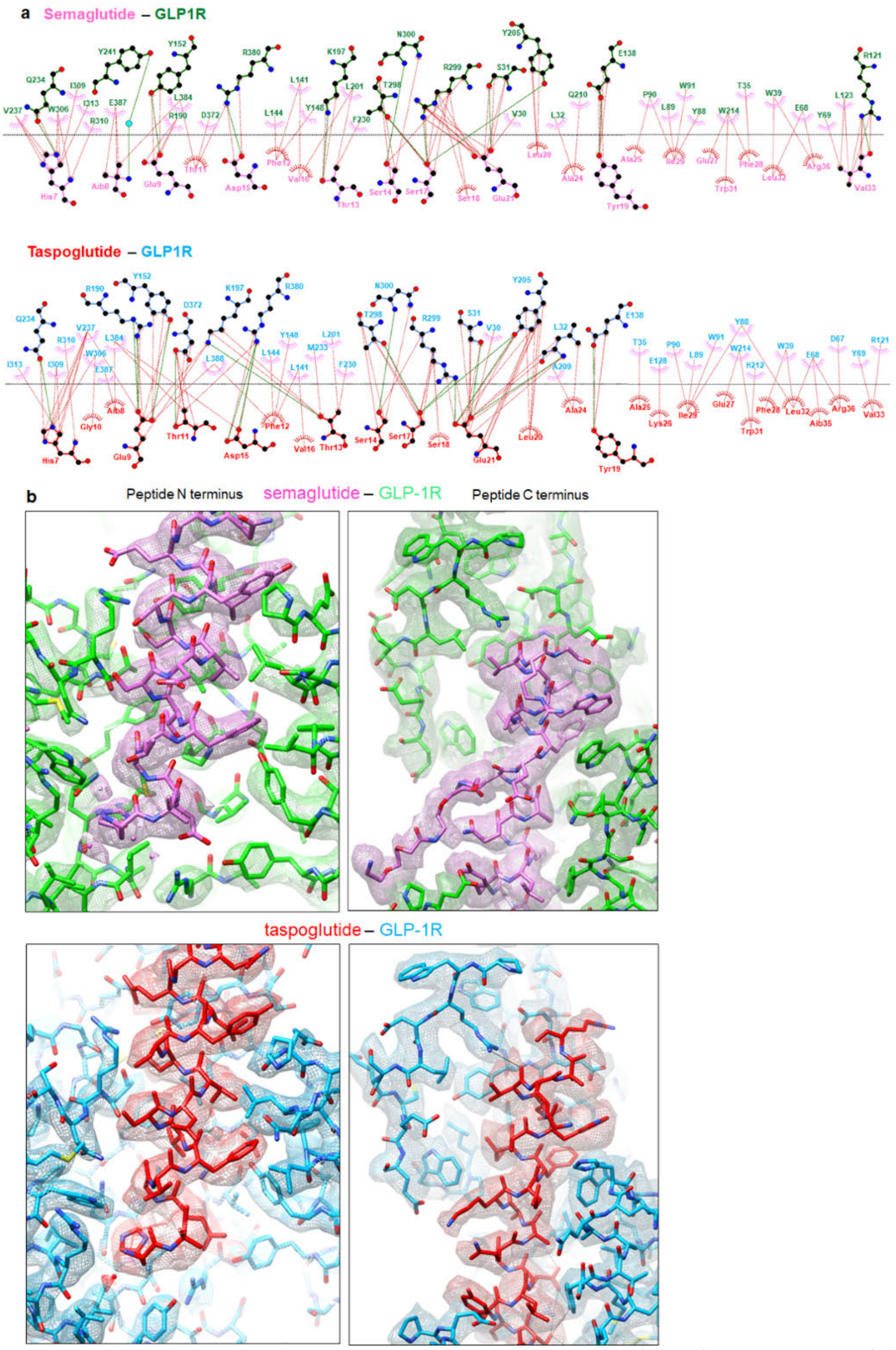


**Figure S4. Ligand interactions with the GLP-1R. a,** Interactions between peptide agonist and GLP-1R as determined by Ligplot+ (Laskowski and Swindells 2011). Top, semaglutide (pink) and the GLP-1R (green); Bottom, taspoglutide (red) and the GLP-1R (blue). GLP-1R residues are located above the dashed black line, and peptide residues below the line. Hydrophobic interactions are illustrated by red (peptide) or pink (GLP-1R) arcs, and interacting residues are joined by a red line. Amino acids involved in H-bonds are shown in atomic detail with H-bonds shown as dashed green lines. **b,** Atomic models of the binding sites of the semaglutide- (upper panels) and taspoglutide- (lower panels) bound GLP-1Rs and their density within the cryo-EM maps. Left, The N-terminal portion of the peptide (residues 7-21) and the GLP-1R TM bundle; Right, C-terminal portion of the peptide (residues 22-36) and the GLP-1R ECD. All residues are displayed as sticks coloured by heteroatom. Colour legends are displayed on the figure panels. *Related to Figures 2 and 3, and Table S2.*

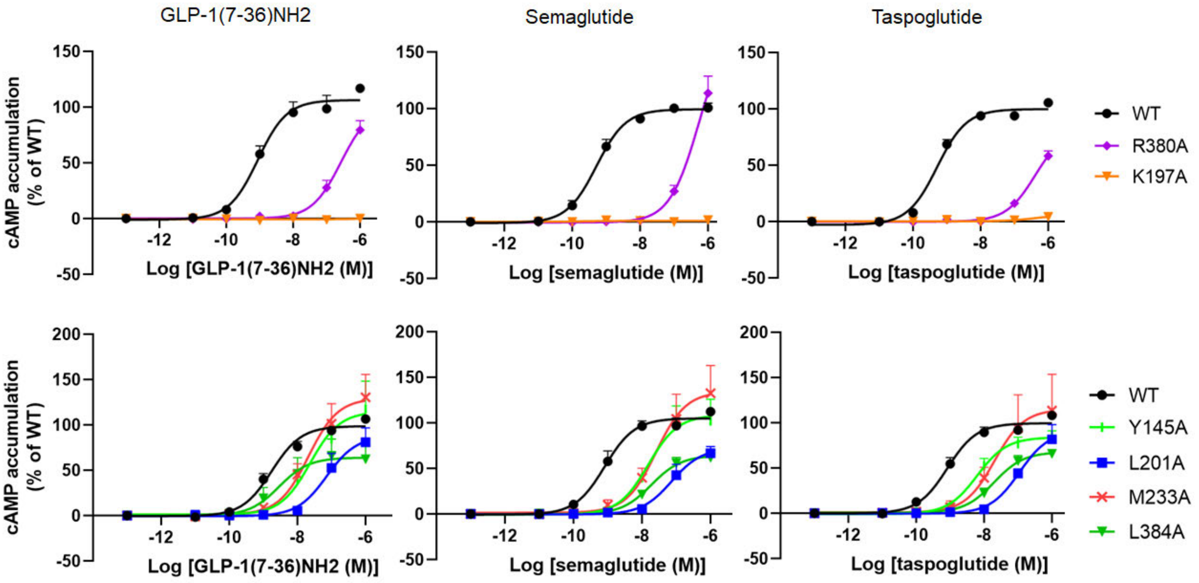


**Figure S5. Ligand mediated cAMP accumulation in ChoFlpIn cells stably expressing WT or mutant hGLP-1Rs.** Concentration response curves for alanine mutants of the GLP-1R were assessed for GLP-1 (left), semaglutide (middle) and taspoglutide (right). Colours identifying the mutant receptors are highlighted on the figure panels. All data are the means + S.E.M. of at least four independent experiments performed in duplicate. *Related to Figures 2 and S4.*

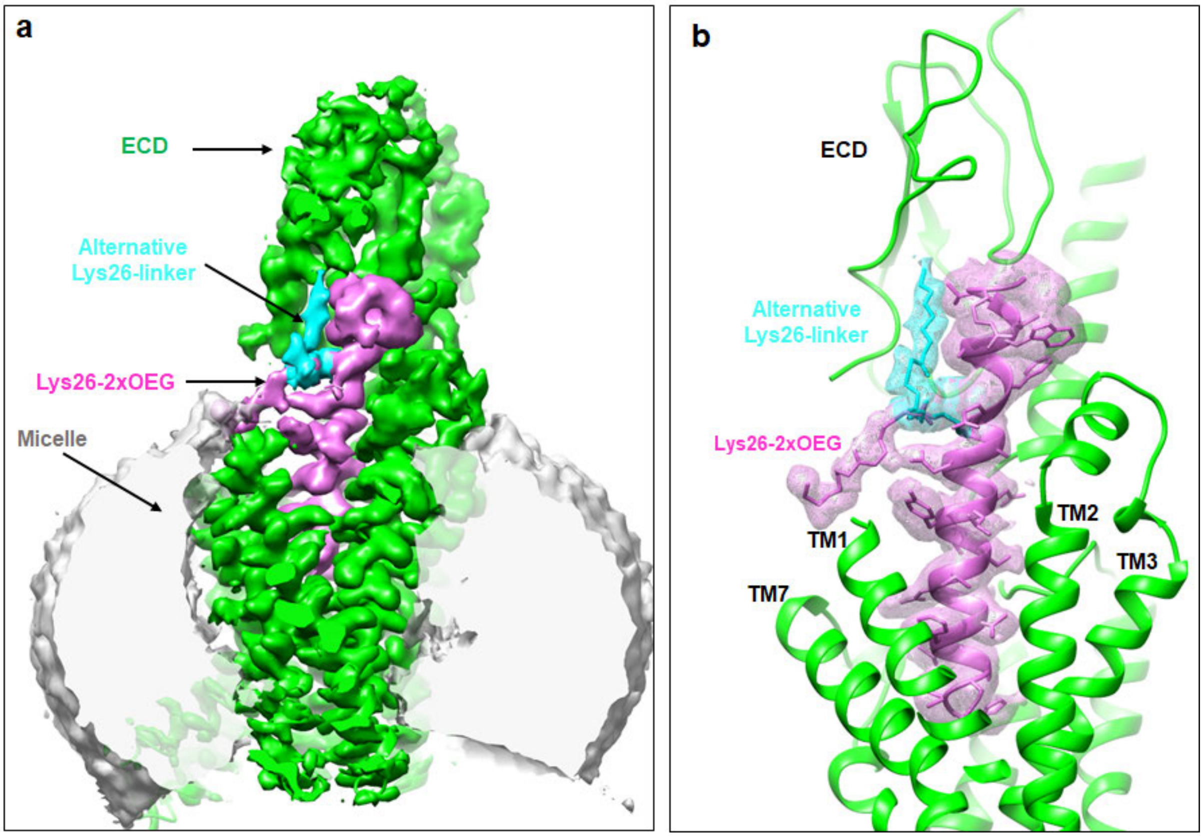


**Figure S6. Model of semaglutide Lys26-2xOEG in the cryo-EM map. a,** Cryo-EM maps (surface) generated from ECD focused map via a zone of 2 Å and mask on semaglutide and the bound GLP-1R structure, with the micelle independently displayed using UCSF chimera. Colours are highlighted on the figure panel. **b**, Model of semaglutide within binding side, illustrating the density for the peptide in the cryo-EM map. Two Lys26-2xOEG models were independently built into the electron density (mesh). These are highlighted in stick form with different colours for the alternate conformations (the major conformation in pink, the alternative one in cyan). The GLP-1R is shown in backbone ribbon format in green. *Related to Figure 3.*

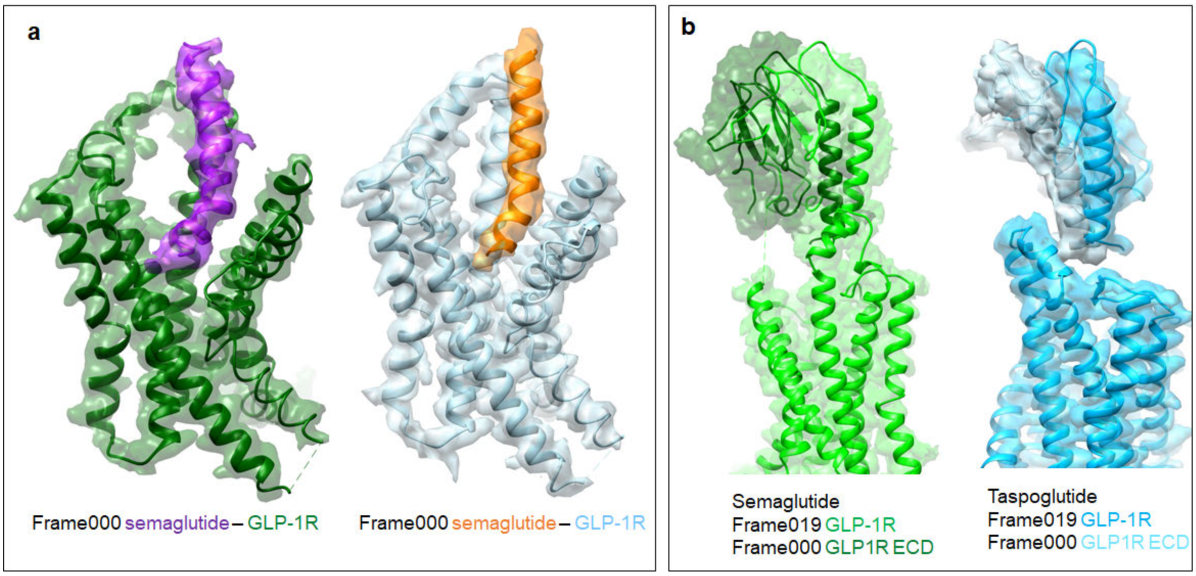


**Figure S7. Orthogonal views of the cryo-EM maps generated in the 3D variability analyses. a,** Orthogonal views of semaglutide- (left) or taspoglutide- (right) bound GLP-1R, displayed in backbone ribbon format, in the cryo-EM maps of the dynamic states (frame 000) from the principal component 1; **b,** Orthogonal views of semaglutide- (left) or taspoglutide- (right) bound GLP-1R in the the cryo-EM maps of frame 000 and frame 019 from the principal component 4. Colours are highlighted on the figure panels. *Related to Figures 5, 6 and S2, and Videos S1 and S2.*
